# Frequent birth of *de novo* genes in the compact yeast genome

**DOI:** 10.1101/575837

**Authors:** William R. Blevins, Jorge Ruiz-Orera, Xavier Messeguer, Bernat Blasco-Moreno, José Luis Villanueva-Cañas, Lorena Espinar, Juana Díez, Lucas B. Carey, M. Mar Albà

## Abstract

Evidence has accumulated that some genes originate directly from previously non-genic sequences, or *de novo*, rather than by the duplication or fusion of existing genes. However, how *de novo* genes emerge and eventually become functional is largely unknown. Here we perform the first study on *de novo* genes that uses transcriptomics data from eleven different yeast species, all grown identically in both rich media and in oxidative stress conditions. The genomes of these species are densely-packed with functional elements, leaving little room for the co-option of genomic sequences into new transcribed loci. Despite this, we find that at least 213 transcripts (~5%) have arisen *de novo* in the past 20 million years of evolution of baker’s yeast-or approximately 10 new transcripts every million years. Nearly half of the total newly expressed sequences are generated from regions in which both DNA strands are used as templates for transcription, explaining the apparent contradiction between the limited ‘empty’ genomic space and high rate of *de novo* gene birth. In addition, we find that 40% of these *de novo* transcripts are actively translated and that at least a fraction of the encoded proteins are likely to be under purifying selection. This study shows that even in very highly compact genomes, *de novo* transcripts are continuously generated and can give rise to new functional protein-coding genes.

## Background

A fascinating mechanism for the formation of new genes known as *de novo* gene birth has gained increased attention in the scientific community (Tautz and Domazet-Lošo 2011; McLysaght and Guerzoni 2015). In contrast to genes formed by gene duplication or gene fusion, *de novo* genes arise from previously non-coding sequences of the genome. Consequently, *de novo* genes can represent veritable leaps of evolutionary innovation. The archetypal version of *de novo* birth begins with a non-genic sequence that undergoes a series of changes which enable it to be transcribed, translated, and potentially confer a new function. While it may seem highly improbable that a few tweaks to noncoding DNA could result in a beneficial novel gene, numerous examples have been described in the literature (Begun et al. 2006; Levine et al. 2006; Li et al. 2010; Toll-Riera et al. 2009; Knowles and McLysaght 2009; Zhuang et al. 2019).

*De novo* gene birth events occurring in eukaryotes are likely to be facilitated by pervasive transcription across the genome, as well as the pervasive translation of the ORFeome, which exposes new loci to selection (Wilson and Masel 2011; Carvunis et al. 2012; Neme and Tautz 2016; Ruiz-Orera et al. 2018, 2014). A small fraction of these ‘accidentally’ transcribed and translated loci could eventually become integrated in cellular pathways and perform essential functions (Chen et al. 2013; Gubala et al. 2017; Xie et al. 2019). The formation of genes with completely new sequences and functions may be key to explaining some of the phenotypic differences between closely-related species (Luis Villanueva-Cañas et al. 2017; Baalsrud et al. 2018; Li et al. 2015).

It has also been hypothesized that *de novo* genes could lead to new biological responses to environmental stress (Arendsee et al. 2014). Prior studies in yeast have supported this idea-they found that a significant fraction of the putative *de novo* genes appeared to be more actively translated in starvation conditions than in rich media (Wilson and Masel 2011; Carvunis et al. 2012). For example, Carvunis et al. 2012 identified over a thousand putative translated ORFs (which they call ‘protogenes’) which are unique to *Saccharomyces cerevisiae*, and approximately one third of them were exclusively translated in specific conditions.

The process of *de novo* gene birth appears to be universal to all organisms. Studies have identified *de novo* genes in several kingdoms; for example, baker’s yeast (Carvunis et al. 2012; Vakirlis et al. 2018), fruit fly (Zhou et al. 2008; Palmieri et al. 2014; Zhao et al. 2014), *Arabidopsis* (Li et al. 2016), and mammals (Toll-Riera et al. 2009; Ruiz-Orera et al. 2015; Guerzoni and McLysaght 2016; Schmitz et al. 2018; Wilson et al. 2017; Wu et al. 2011). A common feature that is shared between these studies is that *de novo* genes typically encode small proteins, the coding sequences tend to include infrequent codons, and they often display patterns of rapid evolution. Less is known about how *de novo* genes appear in the first place; in mammals there is evidence that new transcripts can emerge from the large sequence space of intergenic and intronic genomic sequences via the formation of new signals for the initiation and termination of transcription (Ruiz-Orera et al. 2015). In other groups, such as the Saccharomycotina yeast, introns and intergenic regions are much smaller which could seemingly limit the chances for *de novo* gene birth events. Despite this, previous studies using ribosome profiling data have detected hundreds of evolutionary recent events of new translation initiation (Ingolia et al. 2009; Carvunis et al. 2012; Éléonore Durand et al. 2018).

Estimating the rate of *de novo* gene birth is challenging; the prevalence of this mechanism for new gene creation is still a matter of debate (Light et al. 2014; Casola 2018). On the one hand, studies which incorporate expression data (RNA-Seq and/or Ribo-Seq) find evidence of numerous new transcriptional events, and that many of the new transcripts are likely to be translated (Carvunis et al. 2012; Schmitz et al. 2018; Ruiz-Orera et al. 2014; Lu et al. 2017; Ruiz-Orera et al. 2015). On the other hand, studies based mostly on annotated protein-coding genes and genomic comparisons result in much lower (and more conservative) estimates of *de novo* gene birth rates (Guerzoni and McLysaght 2016; Vakirlis et al. 2018; Ekman and Elofsson 2010). Each approach has limitations-for example, using transcriptomics data for the focal species, but not including the same data for the rest of species, would lead to overestimates of species-specific genes. Similarly, a purely annotation-based analysis is likely to be biased by the varying quality of the annotations of different species. Small proteins (< 100 aa) are often missing in the annotations, and non-model species generally have lower quality reference annotations; this decreases the scope of sequence homology searches. Due to differences in the data and methodologies used by previous studies, groups which attempted to characterize *S. cerevisiae-* specific *de novo* genes have come up with numbers which varied by orders of magnitude-from over a thousand using all predicted ORFs with ribosome profiling data (Carvunis et al. 2012) to a dozen using gene annotations (Vakirlis et al. 2018).

To better understand the dynamics of *de novo* gene emergence in the Saccharomycotina subphylum we designed a strategy that combines gene annotations and transcriptomics data for 11 yeast species. We generated RNA-Seq data for all 11 species which were grown in parallel in two conditions-rich media and oxidative stress. This comparative transcriptomics approach, taken together with an analysis of genomic synteny, led us to identify hundreds of *de novo* transcript candidates in *S. cerevisiae.* We subsequently assessed the coding potential of these *de novo* candidates with our own ribosome profiling data in the same two conditions. Our approach adapts some of the methods used by previous studies and provides new information about the processes underlying the birth of new genes.

## Results

### Assembling novel transcripts from 11 yeast species

We selected 10 species from the Saccharomycotina subphylum including the model organism *S. cerevisiae*, as well as a more distant outgroup species *(Schizosaccharomyces pombe)*, due to their evolutionary history as well as their inclusion in other relevant studies (Figure 1, Supplementary Table 1). All 11 species of yeast were grown in rich media, henceforth referred to as ‘normal’ conditions, as well as in oxidative stress conditions induced by adding hydrogen peroxide (H_2_O_2_) to the rich media, henceforth referred to as ‘stress’.

**Figure 1.**
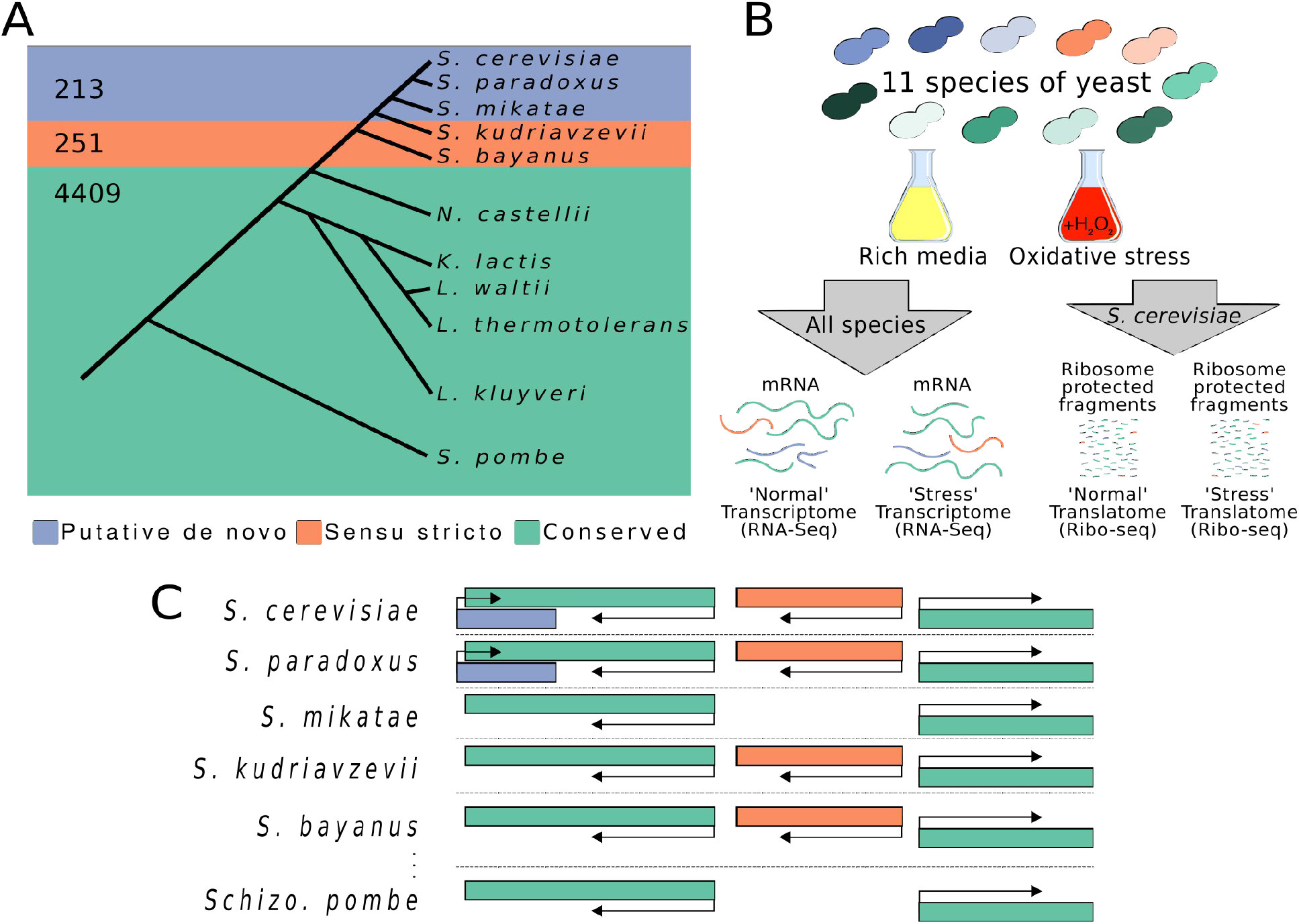
Over two hundred *S. cerevisiae* transcripts are likely to have arisen *de novo* over the past 20 Million years. **A**. We estimated the conservation of each transcript in the yeast phylogeny using homology and synteny. Only transcripts with expression level > 15 TPM, which could be reliably assembled from the RNA sequencing data, were considered. The transcripts were divided in three classes: ‘putative *de novo’* (213 transcripts), *‘sensu stricto’* (251 transcripts) and ‘conserved’ (4,409 transcripts). **B**. Our experiment included 11 species of yeast grown in 2 conditions; rich media and oxidative stress. We generated transcript expression data (RNA-Seq) for all 11 species in both conditions, as well as translational activity data (Ribo-Seq). **C**. Example of transcription on both strands, in a syntenic genomic region, across several species. In green are transcripts which have orthologs in distantly-related species i.e. ‘conserved’, orange are transcripts which have orthologs in either *S. kudriavzevii* or *S. bayanus*, and in purple are transcripts which are only found in *S. cerevisiae, S. paradoxus*, or *S. mikatae.* The putative *de novo* transcript in *S. cerevisiae* and *S. paradoxus* is overlapping a conserved transcript on the opposite strand. The *sensu stricto* transcript is neighboring a conserved transcript which is transcribed in the opposite direction; neighboring divergent transcripts (like this pair) could share a bidirectional promoter.

We performed high throughput RNA sequencing (RNA-Seq) in normal and stress conditions for the 11 species, and assembled *de novo* transcriptomes using Trinity (Grabherr et al. 2013). We combined the set of *de novo* assembled transcripts with the reference annotations to generate an inclusive and non-redundant set of transcripts for each species (Supplementary Figure 1). Our *de novo* assemblies contained an average of 770 novel (unannotated) transcripts for each species studied (Supplementary Table 2).

During the library preparation step we added synthetic spike-in transcripts from the ERCC spike-in kit to each sample. This spike-in allowed us to determine the lower limit of detection of our sequencing pipeline; we could reliably detect if an annotated transcript was present in a given sample for expression values >2 transcripts per million (TPM), but to ensure the complete and reliable *de novo* assembly of a transcript we needed significantly more reads which corresponded to an expression of >15 TPM (Supplementary Figure 2). (Supplementary Table 3; Supplementary Figure 3). As orthologous genes from closely related species generally show similar expression values (McManus et al. 2014), by imposing a cutoff of 15 TPM for all transcripts, including annotated ones, we maximize the likelihood that any existing orthologous transcripts in other species would be recovered by our *de novo* assemblies and thus correctly identified as homologues. Consequently, much of our analysis is focused on transcripts which exceed the stringent cutoff of 15 TPM in either normal, stress, or both conditions. In *S. cerevisiae*, the number of transcripts expressed at >15 TPM amounted to 4,873 transcripts, 385 of which were novel and 4,488 from the reference annotations.

### Identification of *de novo* genes using transcriptomics data, homology searches, and synteny

We used nucleotide and translated nucleotide BLAST homology searches for each transcript, searching for similar sequences in the transcriptomes of each of the 11 yeast species as well as in the proteomes of 35 more distant non-Ascomycota species (Supplementary Table 4). Each species’ BLAST database contained all annotated as well as novel transcripts without any expression cutoff to ensure that the homology search would be as comprehensive as possible. Additionally, we mapped the syntenic genomic regions for the *Saccharomyces sensu stricto* group to detect potential orthologous transcripts whose homology was undetectable with BLAST (Supplementary Table 5). If a transcript had a significant BLAST hit (E-value < 10^−3^) in a transcript/protein from another species, or if it overlapped a transcript in the same genomic position of another species, we considered that the query transcript had a homolog in the other species (Supplementary Table 6). If a homologous transcript was detected in another region of the genome in the query species, we treated this as a paralog-when we classified the conservation of each query transcript, we also considered the conservation of each of the paralogous transcripts.

After performing the homology and synteny searches, we selected the transcripts which were expressed over our threshold (>15 TPM); this amounted to 213 ‘putative *de novo’* transcripts (Supplementary Table 3), including 124 transcripts that had no homologues outside *S. cerevisiae* and 89 that only had homologues in the closely related species *S. paradoxus* or *S. mikatae.* A second group of transcripts, called ‘*sensu stricto’*, had homology in *S. bayanus* and/or *S. kudriavzevii* but not in more distant species (n=251). The last group, which we labeled ‘conserved’, had homologues detected in one or more species outside the ‘*sensu stricto’* group (n=4,409). Although some transcripts in the *sensu stricto* and conserved groups may have also emerged from *de novo* processes, we require more closely related species to trace the step-by-step changes which may result in *de novo* genes.

Perhaps unsurprisingly, the proportion of novel to annotated transcripts was very different in the three groups: The majority of putative *de novo* transcripts were novel (161 out of 213), while the vast majority of conserved transcripts were annotated (4,306 out of 4,406). The *sensu stricto* transcripts were divided into approximately equal parts of annotated and novel transcripts (130 annotated and 121 novel)(Figure 2A, Supplementary Table 7).

**Figure 2.**
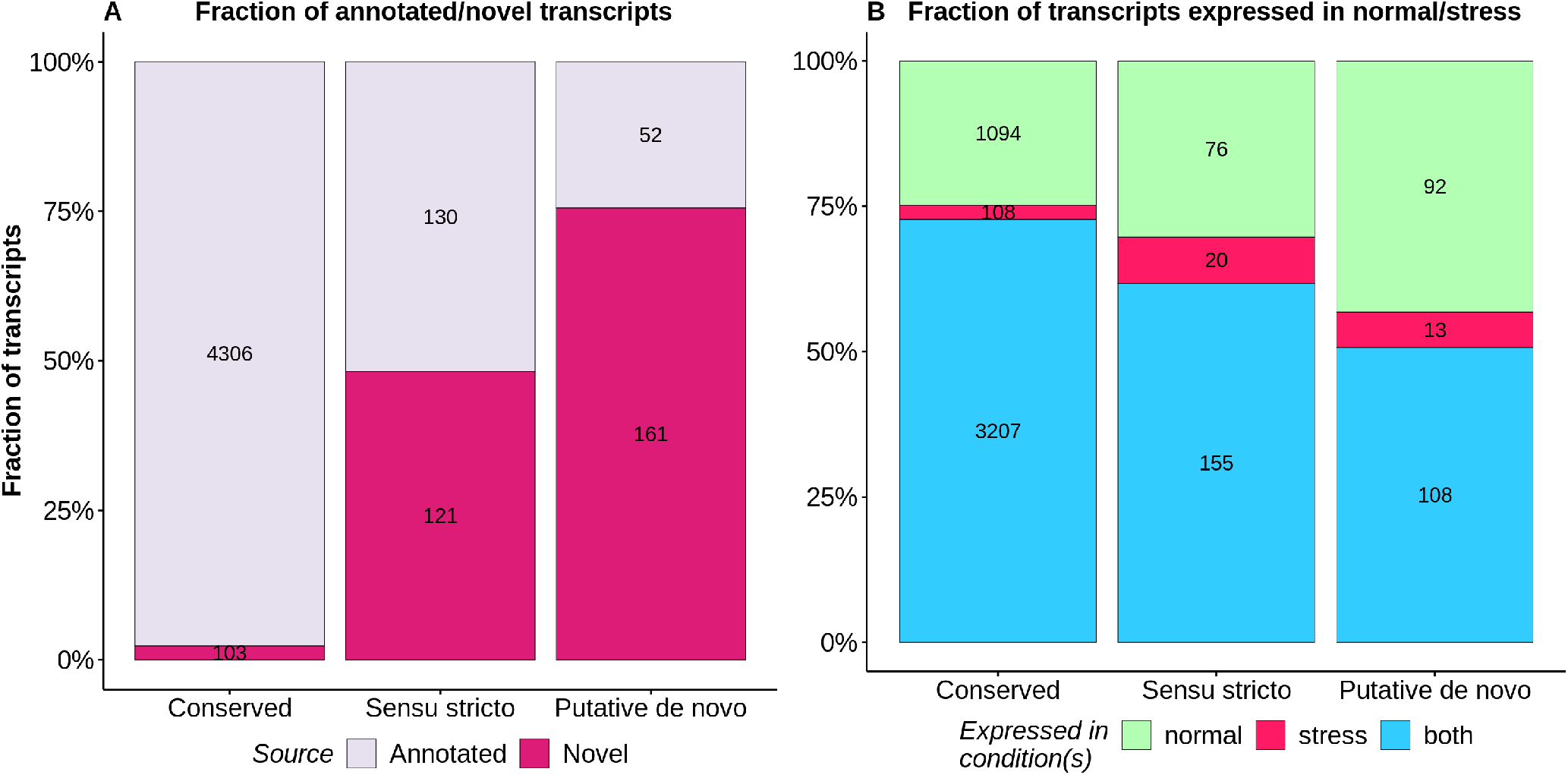
The majority of putative *de novo transcripts* are not annotated and are expressed in both rich media and oxidative stress conditions. **A**. Fraction of transcripts that are annotated (light pink) versus those that are novel (magenta) in the three different conservation levels. Numbers in black text represent the count of transcripts expressed above 15 TPM in each category. **B**. Fraction of transcript expression above 15 TPM in rich media (pale green), in oxidative stress (red), or both conditions (blue). The vast majority of transcripts are either expressed in both conditions. Numbers in black text represent the count of transcripts expressed (above 15 TPM) in each category.

It was previously hypothesized that *de novo* genes may be more expressed in specific conditions (Wu and Knudson 2018; Carvunis et al. 2012). In our dataset, the total number of putative *de novo* transcripts which were expressed in normal conditions was higher than in stress conditions (Figure 2B); only 13 ‘putative *de novo’* transcripts were uniquely expressed in oxidative stress conditions. We may have not been able to assemble some novel transcripts which are only expressed in other conditions not surveyed in this study, but the lower distribution of expression values we observed in the oxidative stress condition (median TPM 7.3 when no expression cutoff is imposed) suggests that their detection and assembly with RNA-Seq could be challenging (Supplementary Table 8).

### Comparing different *de novo* transcript detection strategies

To better understand the impact of using transcriptomics data for multiple species (annotated genes and novel assembled transcripts for 11 species), we ran the same homology and synteny analyses to identify putative *de novo* transcripts, but this time we only used the reference annotations i.e. we excluded novel transcripts. This approach classified 109 annotated *S. cerevisiae* genes as putative *de novo*, compared to 52 with our original approach of reference annotation annotations plus novel transcripts (Supplementary Table 7). The difference corresponds to 5 7 annotated genes (109-52) for which we detected unannotated homologues in other species; therefore the conservation level of these genes would be underestimated if the approach is solely based on the reference annotations. In addition, our transcriptomics data allowed us to detect 161 novel putative *de novo* transcripts that would have been missed by the reference annotations-only approach.

We tested a third possible approach-including transcriptomics data for the focal species, *S. cerevisiae*, in addition to the reference annotations for all species surveyed. In this approach, the number of novel transcripts that were classified as putative *de novo* approximately doubled with respect to our original reference annotations plus novel transcripts approach (336 vs. 161, Supplementary Table 7). This indicates that using transcriptomics data for only the focal species can lead to a considerable overestimation of the number of putative *de novo* transcripts.

These results highlight the importance of including transcriptomics data for many closely-related species when attempting to determine the taxonomic conservation of a given feature. This is especially true for species with incomplete reference annotations, as seems to be the case for some of the species included in this study. Systematic overestimation or underestimation of taxonomic conservation could skew our understanding of the prevalence of *de novo* gene birth, as well as the factors which may contribute to turnover.

### Many *de novo* transcripts overlap other transcripts on the other strand

Approximately 70% of the *S. cerevisiae* genome is spanned by the coding sequences of annotated genes; however, this number is not including untranslated regions, promoters, and other regulatory elements. The relatively high density of this genome leaves little room for new transcriptional events which do not overlap preexisting genes. To test this, we identified all pairs of transcripts which overlapped other transcripts on the opposite strand. We found that there was a clear excess of *de novo* transcripts which had antisense overlap (Figure 3A). Analysis of the spike-in RNA assemblies showed that this was not due to spurious read orientation or other mapping artifacts (Supplementary Figure 6). Additionally, the same excess of overlapping antisense transcripts which was observed in all *de novo* transcripts was also true for the subset of annotated *de novo* genes (Supplementary Table 9).

**Figure 3.**
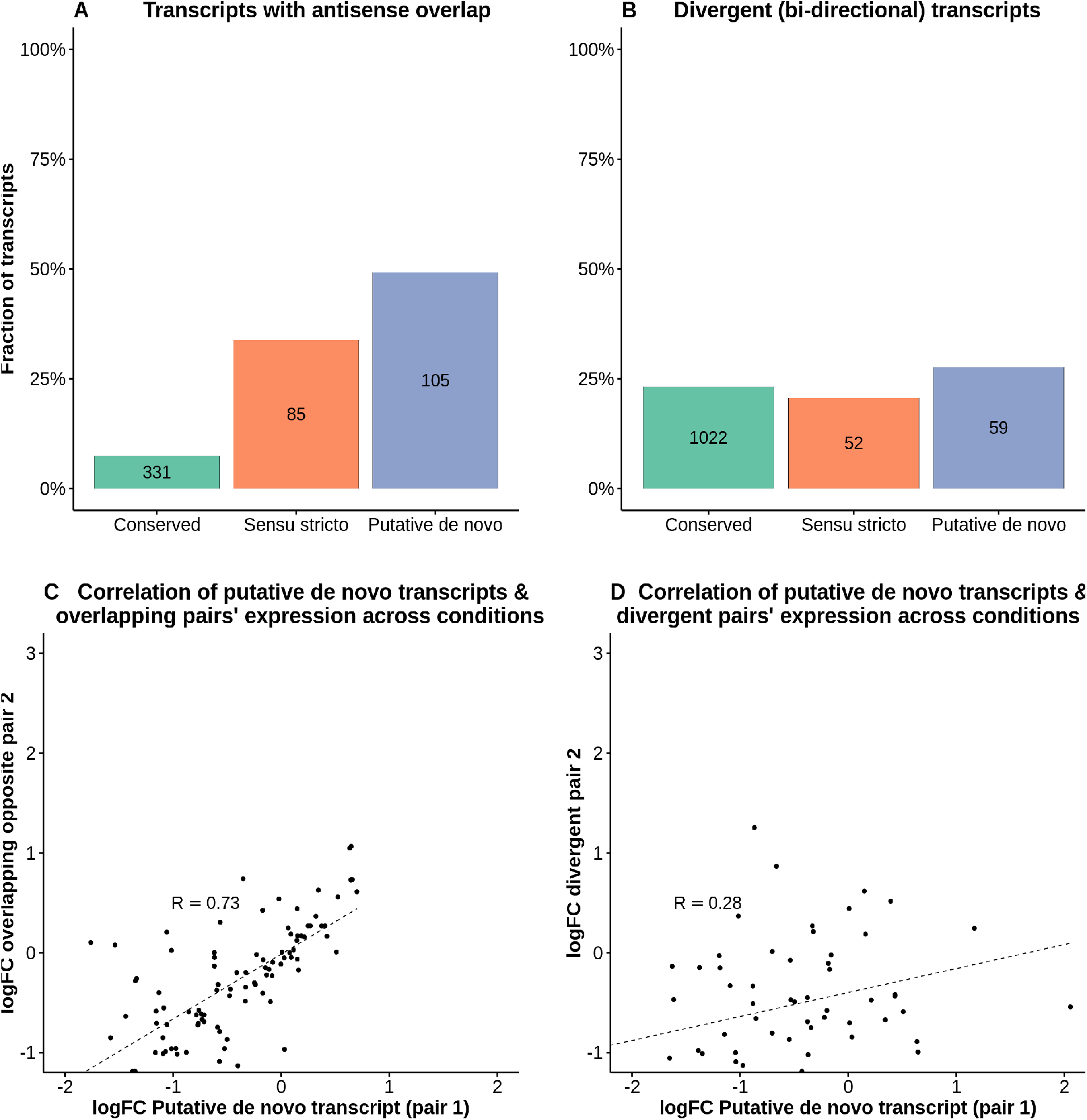
De novo genes often overlap other genes in the opposite orientation. **A**. Fraction of transcripts which overlap genes on the opposite strand. **B**. Fraction of transcripts sharing a bidirectional promoter (divergent). **C**. Comparing the change in expression between conditions of putative *de novo* transcripts and their overlapping antisense pair. ‘logFC’= log(TPM stress/TPM normal). The correlation is highly significant (R= 0.73 Spearman’s correlation, p-value < 10^−5^) D. Comparing the change in expression between conditions of putative de novo transcripts and their divergent (neighboring antisense) pair. The correlation is much weaker (R=0.28 Spearman’s correlation, p-value= 0.02754).

The changes in the expression of the overlapping *de novo* transcripts, measured in normal vs. stress conditions, showed a high correlation with the changes in expression of the other overlapped transcript (Figure 3C, R=0.73, p-value < 10^−5^). In other words, if the relative expression of the sense transcript was higher in stress conditions, the expression of the overlapping antisense transcript also tended to increase, and *vice versa*. A control in which the overlapping pairs were randomized indicated that no correlation was expected by chance (Supplementary Figure 4A). A significant positive correlation was also found in the subset of overlapping pairs that did not include any pairs with putative *de novo* transcripts (Supplementary Figure 4C), indicating that this is a general property of overlapping features.

Next, we examined the degree of overlap between the overlapping gene pairs. In the case of putative *de novo* transcripts, the overlap was very high in most cases with about one third of them overlapping another transcript for the entirety of their length (Supplementary Figure 5). Considering the cumulative sequences of all 213 *de novo* transcripts together, 43.4% of the total length of these transcripts overlapped other transcripts on the opposite strand. Thus, a large fraction of *de novo* transcript emergence takes place on genomic sequences that are already transcribed.

We also analyzed if there was an excess of *de novo* transcripts expressed from bidirectional promoters, as other studies had observed (Vakirlis et al. 2018). We surveyed all pairs of transcripts which were in a divergent orientation and no more than 400nt apart to select for pairs of divergent transcripts which were most likely to share a single nucleosome free region (Huber et al. 2009). However, we did not observe any apparent trend between taxonomic conservation level and the likelihood to be in a divergent orientation to another transcript (Figure 3B); unlike overlapping antisense pairs, changes in expression levels among divergent pairs of transcripts were only weakly correlated (Figure 3D). Similar controls to those performed for overlapping antisense transcripts indicated that this weak correlation is not expected by chance and may be related to shared regulatory elements (Supplementary Figure 4B and 4D).

### Almost half of *de novo* transcripts are translated with a consistent reading frame

To investigate if these new transcripts were potentially being translated, we performed ribosome profiling (Ribo-Seq) for *S. cerevisiae* in both normal and stress conditions. The sequencing of ribosome protected RNA fragments by Ribo-Seq provides a high-resolution snapshot of where ribosomes are bound; this data can be used to distinguish between pervasive ribosomal association, and the 3 nucleotide periodicity indicative of active translation of an open reading frame (ORFs) (Ingolia et al. 2009). After sequencing and mapping, we determined which ORFs were being translated using the program RibORF (Ji et al. 2015); this program is based on the high three nucleotide read periodicity and patterns of coverage which has been observed in actively translated ORFs. We performed this analysis on all possible ORFs, in all 3 frames of a transcript, excluding smaller ORFs which were fully contained inside a longer ORF in the same frame. The vast majority of the ORFs with evidence of translation started with the canonical ATG codon (98.6%), but we could also detect 74 cases in which translation appeared to start at non-canonical start codons (Supplementary Table 10).

As expected, the vast majority of conserved transcripts (97.2%) showed evidence of translation according to RibORF (Figure 4A, Supplementary Figure 7). In the *sensu stricto* and putative *de novo* groups, 48.2% and 37.5% of transcripts were classified as being actively translated (Figure 4A). The percentage of cases identified by RibORF was similar for *de novo* transcripts overlapping other genes in antisense orientation as for non-overlapping transcripts (Supplementary Table 9). Examples of annotated *de novo* genes with ribosome profiling translation included YGL204-C, which encodes an endoplasmic reticulum protein of unknown function, and YOR316C-A, overlapping an ORF encoding a protein that mediates zinc transport into the vacuole.

**Figure 4.**
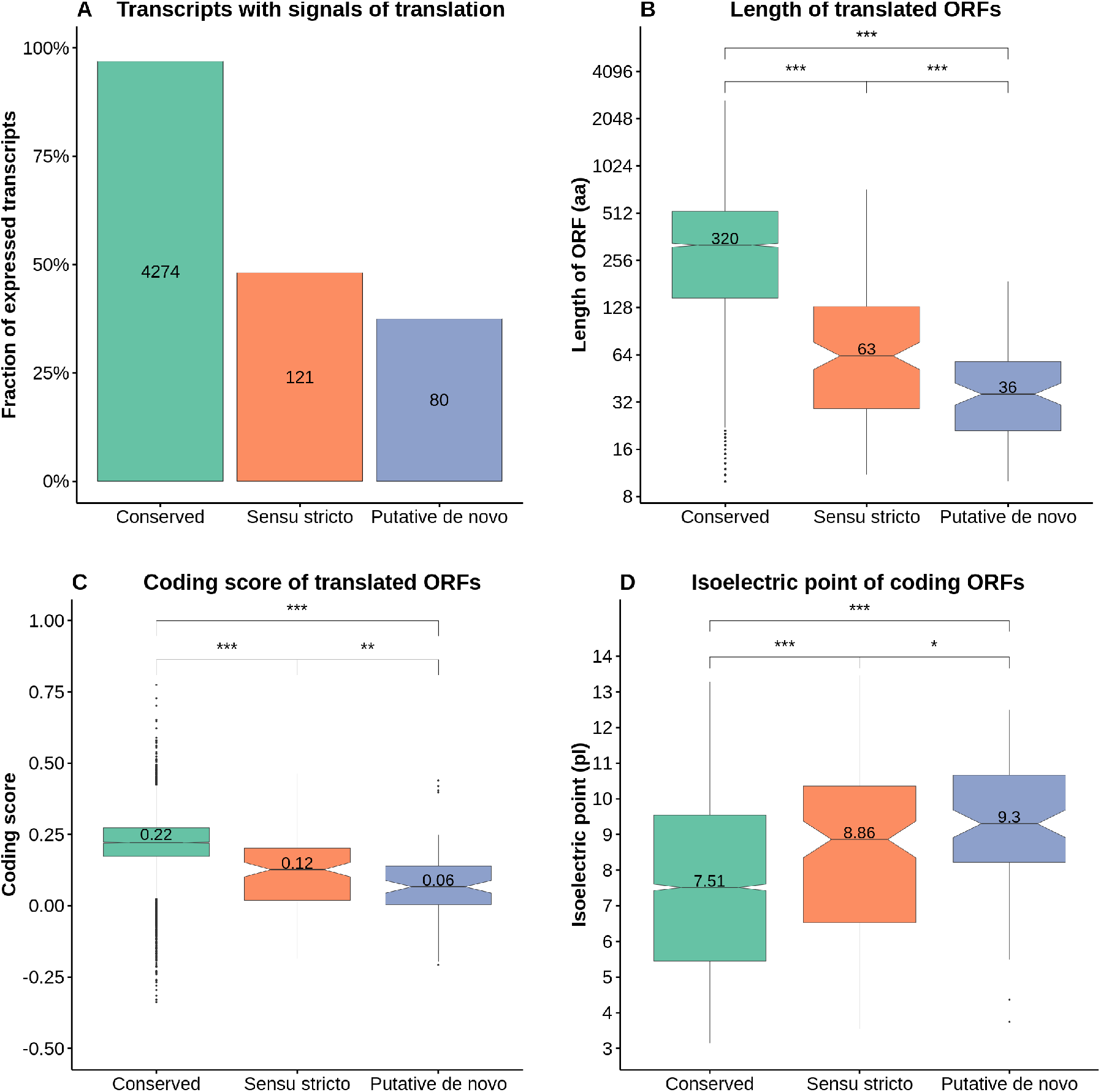
Over 50 putative *de novo* transcripts show evidence of translation. **A.** Identification of transcripts containing putative translated ORFs using ribosome profiling data and annotations. Using ribosome profiling data from yeast grown in rich media and oxidative stress conditions as well as all the annotated CDSs, we identified 80 putative *de novo*, 121 *sensu stricto* and 4274 conserved transcripts with at least one translated open reading frame (ORF). The translated ORFs were detected by RibORF on the basis of high read 3-nucleotide periodicity and uniformity, using a score cutoff of 0.7, in one or both conditions. **B**. Length of translated ORFs. The length of the longest translated ORF per transcript showed a positive relationship with the conservation level. Mean and median ORF length in nucleotides was 960/1142.3 for conserved transcripts, 189/305.2 for *sensu stricto* transcripts and, 108/137.0 for putative *de novo* transcripts. **C** Coding score was calculated using a previously developed hexamer-based metric called CIPHER. Coding score shows a positive relationship with the conservation level. **D** Isoelectric point was predicted with the R package ‘Peptides’, using the EMBOSS pKscale. Significance between the different distributions of conservation levels was calculated with pairwise Wilcoxon tests. *** = p-value < 0.001; ** = p-value < 0.01; * = p-value < 0.05.

The peptides encoded by ORFs in putative *de novo* transcripts tended to be much smaller than those encoded by more conserved genes (Figure 4B), which is in line with previous studies (Toll-Riera et al. 2009; Carvunis et al. 2012; Vakirlis et al. 2018; Tautz and Domazet-Lošo 2011). We measured the coding score in all ORFs with evidence of translation-this metric measures the similarity of the codon composition of the translated ORF and the codon composition of annotated protein-coding sequences (Ruiz-Orera et al. 2015). In general, the coding scores of *de novo* genes were lower than those of more conserved genes (Figure 4C). Finally, we examined if the *de novo* peptides showed abnormally high isoelectric point values, as observed for young mammalian proteins (Luis Villanueva-Cañas et al. 2017). We found that, indeed, the youngest proteins have a higher proportion of basic vs acidic residues (Figure 4D).

Although putative *de novo* transcripts were expressed at lower levels than conserved transcripts (Supplementary Table 7) which could potentially limit the ability of RibORF to correctly identify them as translated, this was not sufficient to explain the lower percentage of translated cases in this group when compared to more conserved genes (Supplementary Figure 8). Therefore, in addition to some protein-coding transcripts, the pool of putative *de novo* transcripts also includes dozens of *bona fide* non-coding transcripts.

### Effect of selection on *de novo* translated ORFs

We searched for signals of purifying selection in translated ORFs using single nucleotide polymorphism (SNP) data from 1,011 *S. cerevisiae* isolates (Peter et al. 2018). We compared the number of observed non-synonymous to synonymous SNPs (PN and PS, respectively) in the different transcript sets to those expected in the absence of purifying selection (Supplementary Table 11); under purifying selection we expect the values of this ratio to be significantly lower than 1 (Ruiz-Orera et al. 2018). In conserved ORFs, the ratio was 0.251, consistent with strong purifying selection (p-value < 10^−3^)(Supplementary Table 11). *Sensu stricto* ORFs showed a value of 0.607 (p-value < 10^−3^), indicating lower but still significant selection. In the case of translated ORFs in putative *de novo* transcripts, only the subset of ORFs with a coding score above the median value in the group (49 ORFs with a coding score > 0.06) showed significant evidence of purifying selection (ratio 0.67, p-value < 0.037). The rest of ORFs in this set of transcripts showed no deviation from neutrality. The data suggest that whereas some of these loci are likely to translate for peptides with biological significance, others may be transiently translated without ever becoming functional.

## Discussion

Comparisons of genes expressed in diverse species point to the existence of many species-or lineage-specific genes with unique sequences, suggesting that many of them may have recently originated *de novo* from previously non-genic sequences (Keese and Gibbs 1992; Albà and Castresana 2005; Neme and Tautz 2013). The birth of new genes can be partially attributed to pervasive transcriptional activity across the genome which continuously produces novel transcripts that could eventually result in new functions (Wilson and Masel 2011; Ruiz-Orera et al. 2014; Neme and Tautz 2016). New promoter sequences arise frequently during the evolution of mammalian genomes (Young et al. 2015) and this is associated with the generation of new expressed loci (Ruiz-Orera et al. 2015). Experiments in *E. coli* have shown that, given an appropriate context, random sequences can act as as promoters after only one mutation or even no mutations (Yona et al. 2018). Here we have generated new data to investigate the evolutionary dynamics of the transcriptomes of several Saccharomycotina yeast, circumventing the biases in previous studies due to differences in the quality of the reference annotations of different species.

Efforts to identify *de novo* protein-coding genes in genomes often use the lack of similar ORFs in the syntenic genomic sequence of closely-related species as additional evidence for the recent origin of the protein (Levine et al. 2006; Cai et al. 2008; Toll-Riera et al. 2009; Knowles and McLysaght 2009).

However, it is not always clear how ORF similarity should be defined; how much shorter can a similar ORF be? Is an ORF located in an equivalent position but showing no evidence of significant sequence conservation a similar ORF? In the case of closely related species such as mouse and rat, conservation of an ORF may occur purely by chance (Ruiz-Orera et al. 2018), which further complicates an ORF-based approach. In the present study we decided to be conservative, so we considered the presence of a transcript in the same syntenic genomic region and strand as an indication of a possible common origin of the two transcripts. We did not consider non-transcribed ORFs, as without transcription, the protein would not be produced.

The lack of annotations of small proteins has strongly hampered any previous attempts to estimate the number of genes that may have arisen *de novo.* Unfortunately, these short and lowly abundant proteins can be particularly difficult to detect with proteomics-based approaches (Slavoff et al. 2013). In order t o obviate this limitation, we generated yeast transcriptomics and ribosome profiling data and performed comparisons across related species that are largely independent of the annotations. This approach is both very sensitive (allowing to detect homologs in other species even if not annotated) and comprehensive (considering all expressed transcripts). The bulk of our analyses focused on *S. cerevisiae* transcripts whose expression was over 15 TPM; this ensured the complete assembly of novel transcripts in our focal species, as well as potentially unannotated orthologs, which permitted us to perform reliable sequence homology searches. Using stringent criteria, we found 213 putative *de novo* transcripts which originated over the last 20 million years. This corresponded to about 4.4% of all *S. cerevisiae* genes expressed at this level (213 out of 4,873); if we do not apply our conservative expression cutoff, this fraction is even higher at 6.2% (436 out of 6,986; Supplementary Tables 3 and 6). BSC4, a previously identified *de novo* gene in *S. cerevisiae (Cai et al. 2008)*, was in our set of putative *de novo* transcripts expressed below 15 TPM. According to our study, the magnitude of *de novo* transcript origination and retention in *S. cerevisiae* is comparable to previous estimates for gene duplication during the same time period (Hahn et al. 2005).

Data accumulated in recent years have shown that there is continuous formation of new transcripts in genomes (Kapusta and Feschotte 2014; Ruiz-Orera et al. 2015). If these transcripts are integrated into cellular functions remains a matter of debate, but it is clear that they provide new opportunities for novel functional genes to arise (Ruiz-Orera et al. 2014). In multicellular organisms with large genomes such as mammals, there is an abundance of intergenic sequence space in which new transcripts could appear. However, Ascomycota yeasts have compact genomes which are densely packed with functional elements, including coding sequences, small RNA genes, and gene expression regulatory sequences (Dujon et al. 2004). Nevertheless, many putative *de novo* transcripts and translated ORFs have been shown to be pervasive throughout this group (Carvunis et al. 2012; Lu et al. 2017). Our findings help explain this contradiction; about half *de novo* transcripts overlap other transcripts, making use of the other genomic strand as a template for transcription.

Previous experiments in yeast have shown that the expression of overlapping antisense transcripts may lead to a reduction in the protein levels of the other transcript (Huber et al. 2016). However, our data do not support this interpretation-rather, there is high positive correlation between the changes in expression of the two transcripts. It may be that the chromatin opening to express one transcript facilitate the expression of the other transcript. Previous analysis of transcript isoform sequencing (TIF-Seq) in *S. cerevisiae* suggested that a large number of novel transcripts could be overlapping conserved genes on the same strand (Lu et al. 2017). The future availability of similar data for other species may help to clarify the evolutionary dynamics of alternative isoforms and how they might contribute to the protein repertoire in Saccharomycotina.

One important remaining open question is: How many putative *de novo* transcripts are functional? We tried to approach this question by examining the distribution of non-synonymous and synonymous SNPs (PN/PS) for transcripts containing ORFs with evidence of translation, and comparing it to the expected values under a lack of purifying selection. Due to the limited number of SNPs in small ORFs, this analysis had to be performed collectively for groups of ORFs. Only a subset of putative *de novo* ORFs (those with relatively high coding scores) were associated with significant purifying selection signatures. These ORFs had a higher PN/PS ratio than conserved ORFs (0.6 and 0.25, respectively); this could mean that they are subject to weaker purifying selection, or that only a fraction of this subset of proteins is functional. Other ORFs did not show any evidence of purifying selection, supporting that there is a pool of peptides that are not functional *per se* but that they could provide raw material for the evolution of new genes. In line with our results, other studies found limited or no evidence of selection in recently evolved *de novo* proteins (Toll-Riera et al. 2009; Durand et al. 2018; Carvunis et al. 2012; Ruiz-Orera et al. 2018; Vakirlis et al. 2018).

The present study provides a picture of *de novo* gene birth at unprecedented resolution. The unique combination of genomics, transcriptomics, and ribosome profiling data indicates that *de novo* gene birth occurs frequently even in compact yeast genomes. About half of the new transcript sequences in *S. cerevisiae* emerge at genomic locations already occupied by other genes on the opposite strand. This configuration does not appear to prevent the translation of small ORFs in the new transcripts, which could potentially confer new functions. Subsequent mRNA-mediated retrotransposition events may generate new copies in other locations, which could continue to evolve free of the constraints imposed by the previously overlapping antisense gene. Future technological developments, including long read RNA sequencing, targeted proteomics, and automated experimental screenings will enable further investigation of the mechanisms driving *de novo* gene birth.

## Methods

### Yeast material

The 11 yeast strains we used in our analysis (Supplementary Table 1) were selected due to their phylogenetic distribution. Several species which are closely related to *S. cerevisiae* were included to facilitate synteny comparisons. A group of more distant and sparsely distributed species was included as well to broaden the scope of the homology searches. Yeast strains were obtained from the labs of both Lucas Carey and Kevin Verstreppen.

### Experimental conditions

We opted for growth conditions that would accommodate many species of yeast (Tsankov et al. 2010); all 11 strains were grown in a custom rich media at 30°C (Supplementary Figure 9). For each species, we selected an isogenic population from streaked plates, then incubated cultures overnight. We used the overnight culture to inoculate two identical 50mL Erlenmeyer flasks containing 20mL of rich media each. After approximately 6 generations of log phase growth, around OD600 of 0.3, we added H_2_O_2_ to one flask to a final concentration of 1.5mM; after 30 minutes, the yeast cells were harvested and frozen from both the stressed and the unstressed flask. We chose a concentration of 1.5mM hydrogen peroxide as we found that this concentration would approximately halve the growth rate for the species included in our study (Supplementary Figure 10); a treatment period of 30 minutes of H_2_O_2_ was selected to capture the greatest variation in expression during stress response (Gasch et al. 2000). For each sample, four 1.5mL centrifuge tubes of cell culture were extracted, centrifuged at 4°C, and then frozen at −80°C. This protocol was slightly modified for the ribosome profiling experiments to account for the increased demand in raw material for the sequencing protocol (see Ribosome profiling section below).

### RNA sequencing

For each sample, 1μg of total RNA underwent poly(A) filtration with streptavidin-coated magnetic nanobeads; ERCC RNA Spike-In Mix (Rna and Consortium 2005) was added prior to library preparation. We ran paired-end 50bp stranded sequencing on the Illumina HiSeq 2500 platform using v4 chemistry. The total number of mapped reads was between 28-38 Million reads per sample (Supplementary Table 12).

### Ribosome profiling

Cultures were grown in 500ml of rich media in 1L Erlenmeyer flasks; we added cyclohexamide (100 μg/ml final concentration) 1 minute prior to harvesting the cells We harvested the yeast cells via vacuum filtration, suspended them in 500 μl of lysis buffer, then flash-froze them with N2(l). For each sample, 2/3 of the harvested cells were reserved for Ribo-Seq and 1/3 for RNA-Seq.

Cells were lysed using the freezer/mill method (SPEX SamplePrep); after preliminary preparations, lysates were treated with RNaseI (Ambion), and subsequently with SUPERaseIn (Ambion). Digested extracts were loaded in 7%-47% sucrose gradients to evaluate the quality of the samples. Monosomal fractions corresponding to digested polysomes were collected; SDS was added to stop any possible RNAse activity, then samples were flash-frozen with N2(l). RNA was isolated from monosomal fractions using the hot acid phenol method. Ribosome-Protected Fragments (RPFs) were selected by isolating RNA fragments of 28-32 nucleotides (nt) using gel electrophoresis. The protocol described in Ingolia et al. 2012 was used to prepare sequencing libraries for both RPFs and fragmented RNA, with minor modifications (Ingolia et al. 2012). Sequencing was performed on the Illumina NextSeq platform.

### Processing of RNA-Seq data

We trimmed the adapters and low quality bases with Trimmomatic (Bolger et al. 2014) with the following parameters:

*ILLUMINACLIP:$illumina_adapters:2:33:20:2:true LEADING:36 TRAILING:32 SLIDINGWINDOW:4:30 MINLEN:35*

We then used Bowtie2 version 2.2.3 with default parameters (Langmead and Salzberg 2012) to map the trimmed RNA-Seq reads to the reference genome. After mapping the reads, we discarded all reads with > 2 mismatches as well as unpaired reads.

### Assembly of transcriptomes

We used Trinity in genome-guided BAM mode (Grabherr et al. 2013) to perform a *de novo* assembly using the following parameters:

*--normalize_max_read_cov 200 --jaccard_clip --genome_guided_max_intron 1002 --min_kmer_cov 2 --max_reads_per_graph 300000 --min_glue 5 --group_pairs_distance 300 --min_contig_length 200*

In this mode Trinity works with mapped reads, but it does not use the reference genome directly to reconstruct the transcripts. We used Transrate (Smith-Unna et al. 2016) to evaluate the quality of each assembly and refined the parameters of Trinity to achieve a high-quality *de novo* assembly. As Trinity does not use the reference genome directly to assemble transfrags, we used GMAP (Wu and Watanabe 2005) to map the assembled transcripts back to the reference genome. We used Cuffmerge from the Cufflinks suite version 2.2.0 (Trapnell et al. 2012) to combine the *de novo* assemblies from normal and stress conditions with the reference transcriptome. When we combined novel and annotated transcripts into a comprehensive transcriptome, novel transcripts from our assembly which overlapped the reference annotations were considered redundant and were excluded from most of the analysis; however, these transcripts were still included in the BLAST database during homology searches.

### BLAST homology searches

The transcripts from each species were subjected to an all-against-all homology search using BLASTn and tBLASTx, not considering matches on the opposite strand, with relaxed e-value of 1e-3. A BLASTx homology search was also performed against the proteomes of 35 distant “outgroup” species (Supplementary Table 4). The results of all BLAST searches were used to create matrices of transcript conservation between species. The BLAST databases contained all annotated as well as novel transcripts from our assemblies, without any expression cutoff. With regards to BLASTn, we only considered hits whose alignment was over 100nt. We used the following parameters:

BLASTn: *-evalue 1e-3 -max_target_seqs 10000*

tBLASTx: *-evalue 1e-3 -max_target_seqs 10000 -strand plus -seg yes*

BLASTx: *-evalue 1e-3 -max_target_seqs 10000 -strand plus -seg yes*

Transcripts which had homologues in most of the 10 other species as well as in the outgroup were considered to be highly conserved, whereas transcripts which had no overlap or similarity to transcripts in any other species were classified as species-specific. Species-specific transcripts, as well as transcripts which are only found in two or three closely-related species, are excellent candidates for tracing the origins of potential *de novo* genes. The results of all homology searches were combined to maximize the detection of homologs in other species; this approach may generate some false positives (spurious homology hits), thus overestimating the conservation of some transcripts, but our objective was to create a set of transcripts with a high likelihood of *de novo* origins.

Homologous transcripts found in the same species were treated as paralogs; we recorded the most distant homology hit for all paralogs of a given transcript. This allowed us to infer potentially deeper conservation for all copies of duplicated genes.

### Genomic synteny comparisons

Syntenic conservation of transcripts was determined with an adapted version of M-GCAT (Treangen and Messeguer 2006), which searches for significant seeds of identical sequences between two genomes called MUMs (maximal unique matches), then sets of parallel, consecutive, and neighboring MUMs are clustered into ‘blocks’ of syntenic segments. Sets of genomic coordinates can then be compared between two species to determine if a transcript in one species overlaps transcripts in the syntenic corresponding region of another species. For each transcript in our focal species (*S. cerevisiae*), we checked if there was conserved synteny in the other *sensu stricto* species; if so, we looked for potential orthologous transcripts on the same strand in the conserved syntenic region which may have eluded BLAST homology searches. If there was no conserved synteny for a given transcript or if there were no transcripts in the syntenic conserved region in the other *sensu stricto* species, we concluded that there were no orthologs according to our synteny comparison.

### Calculating PN/PS

To test if our candidate *de novo* genes were under selective pressure, we analyzed SNPs occurring with a frequency of at least 1% in 1,011 S. cerevisiae isolates (Peter et al. 2018). We classified the observed SNPs as non-synonymous (PN) if the substitutions resulted in the new codon encoding a different amino acid, and synonymous (PS) if they did not. Expected PN and PS values were calculated from SNPs found in intronic regions which did not overlap any features in either strain (Supplementary Table 13); this was the basis for our model of assumed neutral selection. The SNP data were analyzed in groups because the SNP data for single ORFs was not powerful enough to perform statistical tests. We performed a Pearson’s Chi-squared test with Yate’s continuity correction and one degree of freedom to test whether ORFs from varying levels of conservation were under selective pressure (observed PN/PS different from expected PN/PS).

### Prediction of translated ORFs

We used an in-house script to generate genomic coordinates for all possible ORFs for each transcript; this script scans the transcript for canonical and non-canonical start and stop codons, then returns all ORFs which ≥ 3 codons long and are not fully contained in a longer ORF in the same frame. We used RibORF (Ji et al. 2015) to analyze our Ribo-Seq data using the parameters of minimum length=9aa, minimum number of reads=10. RibORF counts the number of reads that fall in each frame and calculates the distribution of reads along the length of the ORF. We used the original proposed cutoff (score ≥ 0.7) to predict translated ORFs-with this cutoff, the vast majority of annotated coding sequences were classified as translated (Supplementary Figure 6).

## Supplementary material and data

Supplementary file contains supplementary tables and figures. Information on the transcripts, including genomic coordinates, expression levels and phylogenetic conservation can be downloaded from https://doi.org/10.6084/m9.figshare.7831259.v1. The raw RNA sequencing data has been uploaded to the SRA under BioProject accession number PRJNA525646, and the raw ribosome profiling data was uploaded under BioProject number PRJNA435567.

## Acknowledgments

We thank Dr. Ksenia Pugach and the Verstreppen lab for cultures of several species of yeast, Leire de Campos-Mata for assistance with the preparation of the RNA for sequencing, and the Sequencing Facilities at the Center for Regulatory Genomics (CRG) and Universitat Pompeu Fabra (UPF).

