## Supplemental Tables and Figures for "Frequent birth of *de novo* genes in the compact yeast genome"

### Supplementary Tables

| Genus | Species | Sample source | Sample Strain | Reference Source | Reference Strain |
|---|---|---|---|---|---|
| *Schizosaccharomyces* | *pombe* | Verstrepen | CBS5682 | NCBI | 972h |
| *Lachancea* | *kluyveri* | Verstrepen | CBS3082 | Genolevures | CBS3082 |
| *Lachancea* | *thermotolerans* | Verstrepen | CHCC5657 | NCBI | CBS6340 |
| *Lachancea* | *waltii* | Verstrepen | CBS6430 | YGOB | Unknown |
| *Kluyveromyces* | *lactis* | Verstrepen | ATCC8585 | NCBI | NRRL Y-1140 |
| *Naumovia* | *castellii* | Verstrepen | CBS4309 | NCBI | CBS4309 |
| *Saccharomyces* | *bayanus* | SCBL | Unknown | SSS | CBS 7001 |
| *Saccharomyces* | *kudriavzevii* | Verstrepen | IFO1802 | SSS | IFO1802 |
| *Saccharomyces* | *mikatae* | Verstrepen | IFO1815 | SSS | IFO1815 |
| *Saccharomyces* | *paradoxus* | SCBL | Unknown | SSS | NRRL Y-17217 |
| *Saccharomyces* | *cerevisiae* | SCBL | S288C | SGD | S288C |

**Supplementary Table 1. Yeast species and strains used for sequencing.** Samples were obtained from the Single Cell Behavior Laboratory (SCBL) collection at the UPF and the Verstrepen Laboratory collection at the VIB. We selected the most complete reference genome and annotations available for each species at the beginning of the project; reference genome and annotation sources include the Saccharomyces genome database (SGD), SaccharomycesSensuStricto.org (SSS), the National Center for Biotechnology Information (NCBI), the Yeast Gene Order Browser (YGOB), and the Genomic Exploration of the Hemiascomycete Yeasts (Génolevures).

| Species | Assemblies | Novel | Gene annotations | Total transcripts |
|---|---|---|---|---|
| *S. cerevisiae* | 6476 | 697 | 6291 | 6988 |
| *S. paradoxus* | 6534 | 630 | 6200 | 6830 |
| *S. mikatae* | 6613 | 730 | 5998 | 6728 |
| *S. kudriavzevii* | 6441 | 722 | 5892 | 6614 |
| *S. bayanus* | 6522 | 432 | 5995 | 6427 |
| *N. castellii* | 5938 | 267 | 5871 | 6138 |
| *K. lactis* | 6056 | 929 | 5413 | 6342 |
| *L. waltii* | 6185 | 1215 | 5524 | 6739 |
| *L. thermotolerans* | 6220 | 1253 | 5499 | 6752 |
| *L. kluyveri* | 6468 | 868 | 6117 | 6985 |
| *Schizo. pombe* | 7033 | 413 | 6869 | 7282 |

**Supplementary Table 2. Number of transcripts per species.** Here are the descriptions of each row: 'Assemblies'= number of total transcripts in the *de novo* transcriptome assemblies generated with Trinity. 'Novel'= number of transcripts from the *de novo* assemblies which did not overlap any annotated features on the same strand (determined by Cuffmerge). 'Gene annotations'= number of total annotated transcripts in each species. 'Total transcripts'= number of novel transcripts plus annotated transcripts. No expression cut-off was used in this table, but the parameters used in the de novo assembly (Trinity) required a minimum coverage of RNA-Seq reads when evaluating each potential transfrag.

| Cut-off | Conserved | *Sensu stricto* | Putative de novo | All |
|---|---|---|---|---|
| No minimum | 6143 | 407 | 436 | 6986 |
| > 2 TPM | 5906 | 390 | 416 | 6712 |
| > 15 TPM | 4409 | 251 | 213 | 4873 |

**Supplementary Table 3. Number of different transcripts in *S. cerevisiae* after applying expression cut-offs.** 'TPM'= transcripts per Million. 2 TPM is the lower limit of detection of our pipeline for annotated features as established by the ERCC spike-in kit, and 15 TPM reflects the lower limit for which we are able to fully reconstruct the transcript in our de novo assembly (Supplementary Figure 2). We use 15 TPM as our threshold for all transcripts in the rest of our analysis as we want to ensure that we could fully assembly orthologous transcripts that may not be annotated in other species, even if they are annotated in the focal species. This decision was based on observations that there is a general correlation between the expression levels of orthologous genes.

| **Animal** | **Plant** | **Bacteria** | **Fungi (Non-ascomycota)** | **Protist** |
|---|---|---|---|---|
| *Homo sapiens* | *Arabidopsis thaliana* | *Escherichia coli* | *Parasitella parasitica* | *Entamoeba invadens* |
| *Mus musculus* | *Zea mays* | *Paraburkholderia fungorum* | *Magnaporthe oryzae* | *Plasmodium falciparum* |
| *Gallus gallus* | *Oryza sativa* | *Bacillus subtilis* | *Zymoseptoria tritici* | *Leishmania major* |
| *Xenopus tropicalis* | *Physcomitrella patens* | *Nostoc punctiforme* | *Cryptococcus neoformans* | *Paramecium tetraurelia* |
| *Danio rerio* | *Triticum aestivum* | *Burkholderia multivorans* | *Ustilago maydis* | *Pythium irregulare* |
| *Drosophila melanogaster* | | *Actinobacteria bacterium* | *Puccinia graminis* | |
| *Caenorhabditis elegans* | | *Betaproteobacteria bacterium* | *Rhizoctonia solani* | |
| *Nematostella vectensis* | | | *Allomyces macrogynus* | |
| *Daphnia pulex* | | | *Mitosporidium daphniae* | |

**Supplementary Table 4. Outgroup species for sequence similarity searches.** The proteomes of the 35 listed species were downloaded from Ensembl and concatenated into one file. BLASTX was used to search each yeast transcript against the outgroup proteome database using an e-value cutoff of 0.001.

| | *S. paradoxus* | *S. mikatae* | *S. kudriavzevii* | *S. bayanus* |
|---|---|---|---|---|
| *S. cerevisiae* | *91* | *86* | *86* | *81* |
| *S. paradoxus* | *100* | *89* | *88* | *83* |
| *S. mikatae* | | *100* | *86* | *80* |
| *S. kudriavzevii* | | | *100* | *87* |

**Supplementary Table 5. percentage of the genome covered by pairwise genomic alignments.** The syntenic alignments were produced with M-GCAT. The values correspond to the percentage of total genome sequence from both species taken together in the alignment. See Supplementary Table 1 for more details on the reference genomes and annotations. As expected, pairs of species which had diverged more recently had larger fractions of their genomes covered by syntenic blocks.

| | Outgr | Schizo pombe | L. kluy L. therm L. waltii K. lactis | N. cast | S. baya | S. kudr | S. mika | S. para | S. cere-specific |
|---|---|---|---|---|---|---|---|---|---|
| | | | **Conserved** | | **Sensu stricto** | | | **Putative de novo** | |
| 1. Transcriptomes | 4605 | 121 | 1361 | 48 | 194 | 90 | 88 | 114 | 363 |
| 2. Genomic synteny | 4605 | 121 | 1361 | 48 | 251 | 158 | 107 | 78 | 257 |
| 3. Paralogs | 4613 | 136 | 1346 | 48 | 247 | 160 | 106 | 73 | 257 |

**Supplementary Table 6. Determining the phylogenetic conservation of *S. cerevisiae* transcripts.** Homology detection was performed in three steps: '1. Transcriptomes'= BLAST-based searches across species at the nucleotide and protein levels. '2. Genomic synteny'= Search for expressed sequences in regions of conserved genomic synteny in the 5 *sensu stricto species*. '3. Paralogs'= First, we performed BLAST-based searches to detect paralogs of a given transcript in the same genome, then we recorded if any paralogs had more distant homologs in other species than the target transcript. 'Distant species'= If homology hits were detected by BLAST in the proteomes of the 35 outgroup species (Supplementary Table 4). 'Outgr' is the number of transcripts with homology hits in outgroup species. In columns from '*Schizo. pombe'* through '*S. paradoxus',* the number represent transcripts with homology hits in the indicated species but not in more distant species. '*L. Kluy / L. therm / L. waltii / K. lactis'*= homology hits to at least one of the species in *Lachancea* or *K. lactis*. '*S. cere*-specific'= transcripts without homologs in any other species. See Figure 1 for complete species names and phylogenetic tree. We discarded any transcript that did not produce a hit against itself in the BLAST searches. The transcripts were classified into three groups, 'Conserved', 'S*ensu stricto'* or 'Putative *de novo',* depending on the depth of conservation. No expression cut-off is applied in this table; see Supplementary Table 3 to see the distribution of transcripts' conservation after applying expression cut-offs.

| Group | | Transcripts per Million (TPM) | | |
|---|---|---|---|---|
| | | **Average** | **Median** | **SD** |
| Conserved | Normal | 148.4 | 29.9 | 554.5 |
| | Oxidative stress | 144.3 | 17.3 | 671.5 |
| *Sensu stricto* | Normal | 136.0 | 17.6 | 1014.0 |
| | Oxidative stress | 156.3 | 13.0 | 1378.5 |
| Putative *de novo* | Normal | 50.1 | 13.0 | 476.2 |
| | Oxidative stress | 46.8 | 7.3 | 511.6 |

**Supplementary Table 7. Descriptive statistics of TPM distribution for different groups of transcripts.** 'Conserved' transcripts are expressed at higher levels than the other two groups. The median expression level of transcripts in oxidative stress conditions was lower than in normal conditions (regardless of conservation), but the standard deviation was greater. No gene expression cut-off was applied in this table.

| Approach | Gene type | Conserved | *Sensu stricto* | Putative *de novo* |
|---|---|---|---|---|
| 1. Transcriptomics and annotations in all species | Novel | 103 | 121 | 161 |
| | Annotated | 4306 | 130 | 52 |
| 2. Gene annotations only for all species | Novel | N/A | N/A | N/A |
| | Annotated | 4303 | 74 | 109 |
| 3. Transcriptomics in reference species only | Novel | 43 | 6 | 336 |
| | Annotated | 4335 | 54 | 97 |

**Supplementary Table 8. Effect of using transcriptomics data *vs.* reference annotations in the classification of transcripts into different conservation levels.** '1. Transcriptomics and annotations in all species'= the RNA-Seq-based approach used in the present study which combines *de novo* transcript assemblies with the reference annotations for each of the species surveyed. '2. Gene annotations only for all species'= the result of using the available gene annotations for all species comparisons. '3. Transcriptomics in reference species only'= the result of using transcriptomics data and reference annotations for the focal species, but only using the reference annotations for the rest of the species. Numbers reflect only those transcripts expressed >15 TPM in at least one condition (normal/stress).

| **Putative *de novo* (213)** | **Antisense overlapping** | | **Not overlapping** | |
|---|---|---|---|---|
| | Translated | Not translated | Translated | Not translated |
| Novel | 25 | 64 | 30 | 42 |
| Annotated | 2 | 14 | 5 | 31 |
| ***Sensu stricto* (251)** | **Antisense overlapping** | | **Not overlapping** | |
| | Translated | Not translated | Translated | Not translated |
| Novel | 26 | 46 | 19 | 30 |
| Annotated | 8 | 5 | 74 | 43 |
| **Conserved (4409)** | **Antisense overlapping** | | **Not overlapping** | |
| | Translated | Not translated | Translated | Not translated |
| Novel | 32 | 54 | 12 | 5 |
| Annotated | 241 | 4 | 3985 | 76 |

**Supplementary Table 9. Number of genes in overlapping antisense pairs, according to translation, source, and conservation group.** 'Anti sense overlapping' was defined as two transcripts on opposite strands which had at least 1 nucleotide overlapping between them. 'Novel'= transcripts which came from our *de novo* assembly using Trinity that were not redundant with annotated features on the same strand. 'Annotated'= transcripts from the reference annotations. 'Translated' = transcripts containing at least one ORF which surpassed our RibORF cutoff of 0.7 in at least one condition, according to our ribosome profiling data.

| **Group (ORFs)** | **Translated with ATG** | **Translated with other NTG** | **Total translated ORFs** |
|---|---|---|---|
| Conserved | 5044 | 35 | 5079 |
| *Sensu stricto* | 122 | 16 | 138 |
| Putative *de novo* | 84 | 13 | 97 |

**Supplementary Table 10. Identification of translated ORF using ribosome profiling data.** We used RibORF with a score cut-off of 0.7 to identify translated ORFs from a set of all predicted ORFs (excluding smaller ORFs which were fully contained in another ORF in the same frame), according to our ribosome profiling data in normal and oxidative stress conditions. 'Translated with ATG'= translated ORFs with canonical ATG codon of size 3 amino acids or longer. To facilitate further analyses on purifying selection, these numbers include several annotated coding sequences that were not detected by RibORF in our samples (26 Conserved, 23 *Sensu stricto*, 35 Putative *de novo*). 'Translated with other NTG'= translated ORFs with non-canonical start sites like CTG/GTG/TTG that did not overlap translated ORFs with ATG sites in the same frame.

| Group (ORFs) | Observed Pn | Observed Ps | Expected Pn | Expected Ps | Pn/Ps norm. (obs/exp) | p-value |
|---|---|---|---|---|---|---|
| Conserved | 51,364 | 80,449 | 94,622 | 37,194 | 0.2510 | $< 10^{-5}$ |
| *Sensu stricto* | 876 | 559 | 1,038 | 399 | 0.6024 | < 10-$^{5}$ |
| Putative *de novo* high coding score | 175 | 102 | 199 | 78 | 0.6724 | 0.037 |
| Putative *de novo* low coding score | 162 | 63 | 163 | 64 | 1.0096 | 1 |

**Supplementary Table 11. Non-synonymous and synonymous single nucleotide polymorphisms (SNPs).** 'Observed PN'= number of observed non-synonymous SNPs with a frequency > 1% in 1,011 different *S. cerevisiae* isolates. 'Observed PS'= number of observed synonymous SNPs with a frequency > 1% in 1,011 different *S. cerevisiae* isolates. 'Expected Pn' and 'Expected Ps' were obtained for each set of sequences using a table of nucleotide substitutions estimated from SNPs in yeast intronic regions (Supplementary Table 13). 'Putative *de novo* high coding score' refers translated ORFs in putative *de novo* transcripts with a coding score ≥ 0.06184 (49 ORFs). 'Putative *de novo* low coding score'= the set of remaining 50 translated ORFs with lower coding scores. 'p-value' refers to the comparison of observed vs. expected Pn / Ps frequencies using a chi-square test.

| species | # mapped reads | # paired reads | % paired reads after filtering and mapping |
|---|---|---|---|
| *S. cerevisiae* | 35290404 | 25995531 | 67.88% |
| *S. paradoxus* | 38148823 | 27820909 | 68.56% |
| *S. mikatae* | 36107601 | 25791495 | 70% |
| *S. kudriavzevii* | 28896378 | 21631444 | 66.79% |
| *S. bayanus* | 23177427 | 22974267 | 50.44% |
| *N. castellii* | 38062640 | 26769999 | 71.09% |
| *K. lactis* | 37346553 | 26293324 | 71.02% |
| *L. waltii* | 31787148 | 24231048 | 65.59% |
| *L. thermotolerans* | 33425642 | 24925275 | 67.05% |
| *L. kluyveri* | 33494394 | 24399133 | 68.64% |
| *Schizo. pombe* | 30184621 | 22389887 | 67.41% |

**Supplementary Table 12. Number of mapped RNA-Seq reads per sample in normal conditions.** '# mapped reads'= the total number of reads which passed our quality control filtering and were mapped by Bowtie2 to the reference genome. '# paired reads' = number of read pairs before quality control and mapping.

| From/to | A | C | G | T |
|---|---|---|---|---|
| **A** | NA | 77 | 385 | 105 |
| **C** | 68 | NA | 62 | 362 |
| **G** | 370 | 55 | NA | 65 |
| **T** | 126 | 373 | 62 | NA |

**Supplementary table 13. Frequency of single nucleotide polymorphisms (SNPs) in introns from *Saccharomyces cerevisiae*.** The SNP data were calculated using annotated introns and SNP data from a study of 1,011 Saccharomyces cerevisiae isolates. Only SNPs with a minimum allele frequency of 0.01 were considered.

### Supplementary Figures

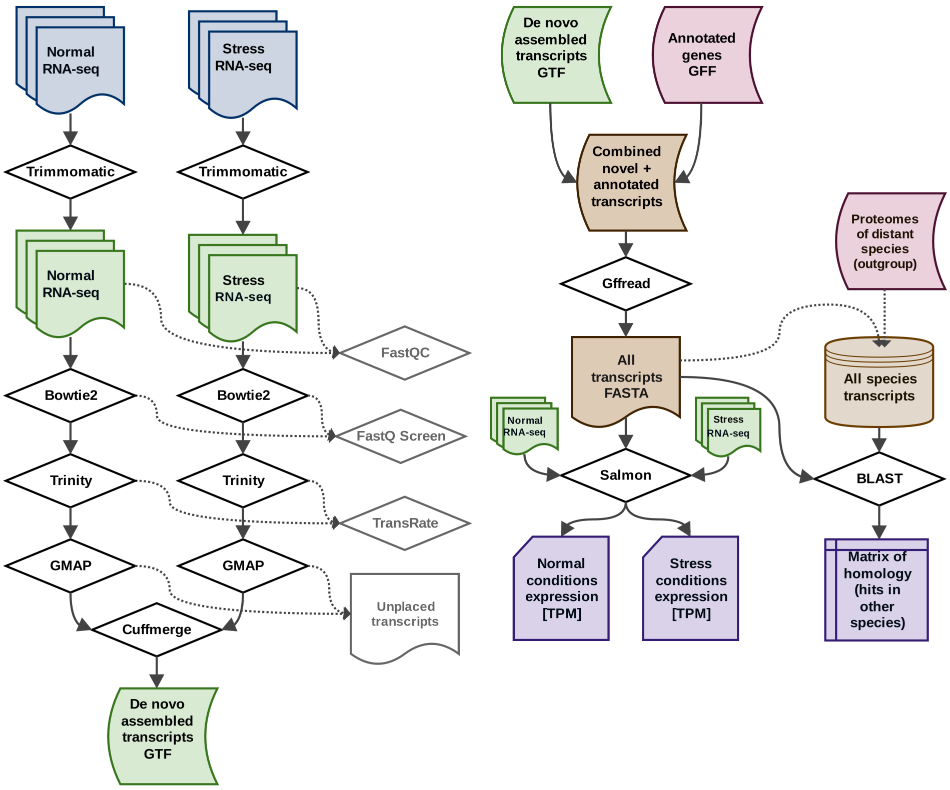


**Supplementary Figure 1. Flow chart of our RNAseq analysis pipeline.** We began our analysis with raw RNA-Seq sequencing fastq files for each of the species both conditions. Adapters and low-quality reads were removed with Trimmomatic, then FastQC was used to do a subsequent quality assessment. The high-quality reads were then mapped to the reference genome with Bowtie2. Trinity was run in reference-free mode, so the assembled transcripts it produces are lacking genomic coordinates. For this reason we used GMAP to map where the assembled transcripts belong on the reference genome. We then used Cuffmerge to compare and combine the reference annotations with our *de novo* assembly. Nucleotide sequences were extracted for each transcript using the tool gffread from the Cufflinks suite, and BLAST databases were created for each species using the complete transcriptome (novel transcripts & annotated transcripts). Each transcript was used as a query in BLAST searches against all BLAST databases (the transcriptomes of all 11 species) as well as the proteomes of 35 distant non-Ascomycota species. Salmon was used to quantify the expression of each transcript in both conditions.

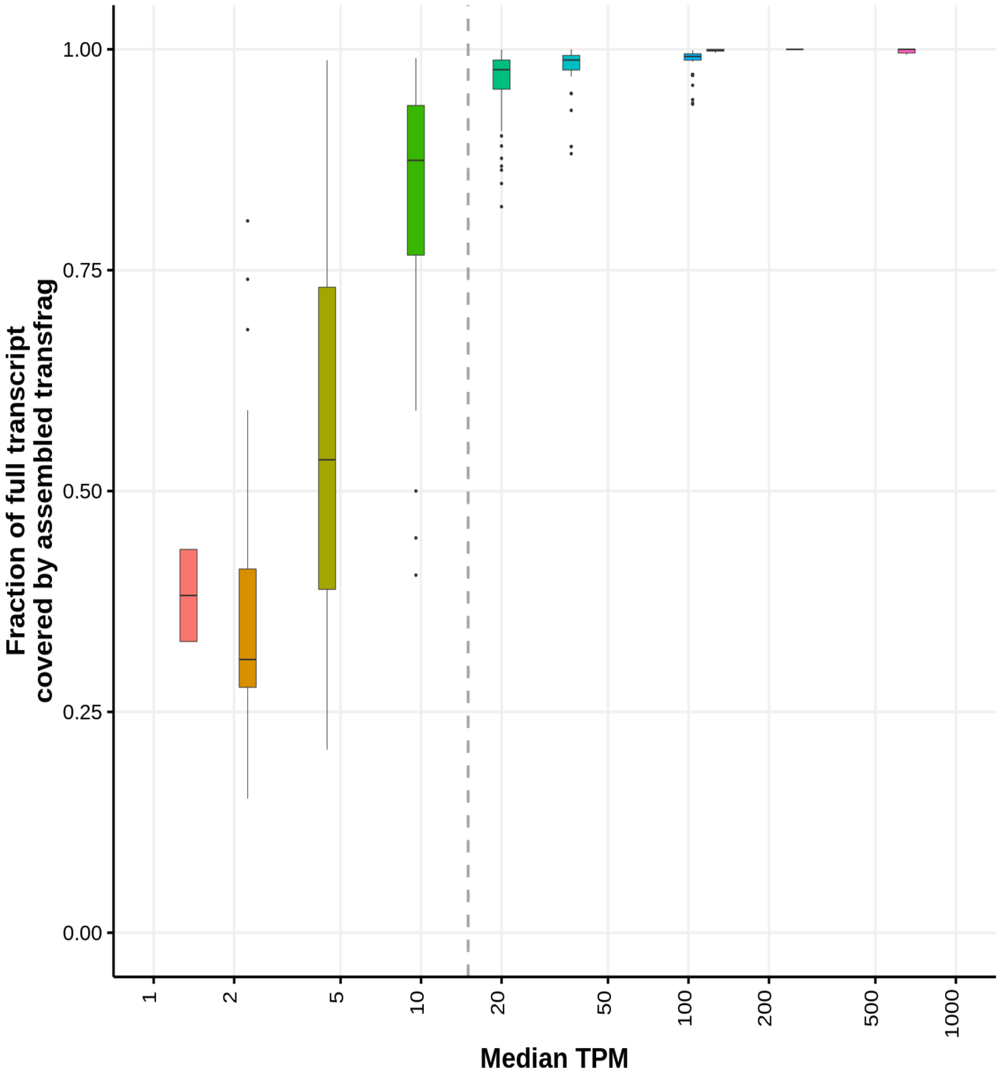


**Supplementary Figure 2. Assessing the relationships between transcript coverage (TPM) and the fraction of transcripts that is fully assembled.** 'TPM'= transcripts per million. We attempted to assemble transcripts for each of the 92 synthetic ERCC spike-in transcripts in each of the 24 samples that we sequenced. As these synthetic transcripts come in different abundances, we estimated the minimum abundance necessary to consistently fully assemble the synthetic transcript with Trinity. Each colored box plot contains the subset of synthetic transcripts at each abundance level; the span of each box represents the range of 'completeness' of the assembly for a given concentration of transcripts. We determined the lower limit of our pipeline to fully and consistently reconstruct a novel transcript to be approximately 15 TPM.

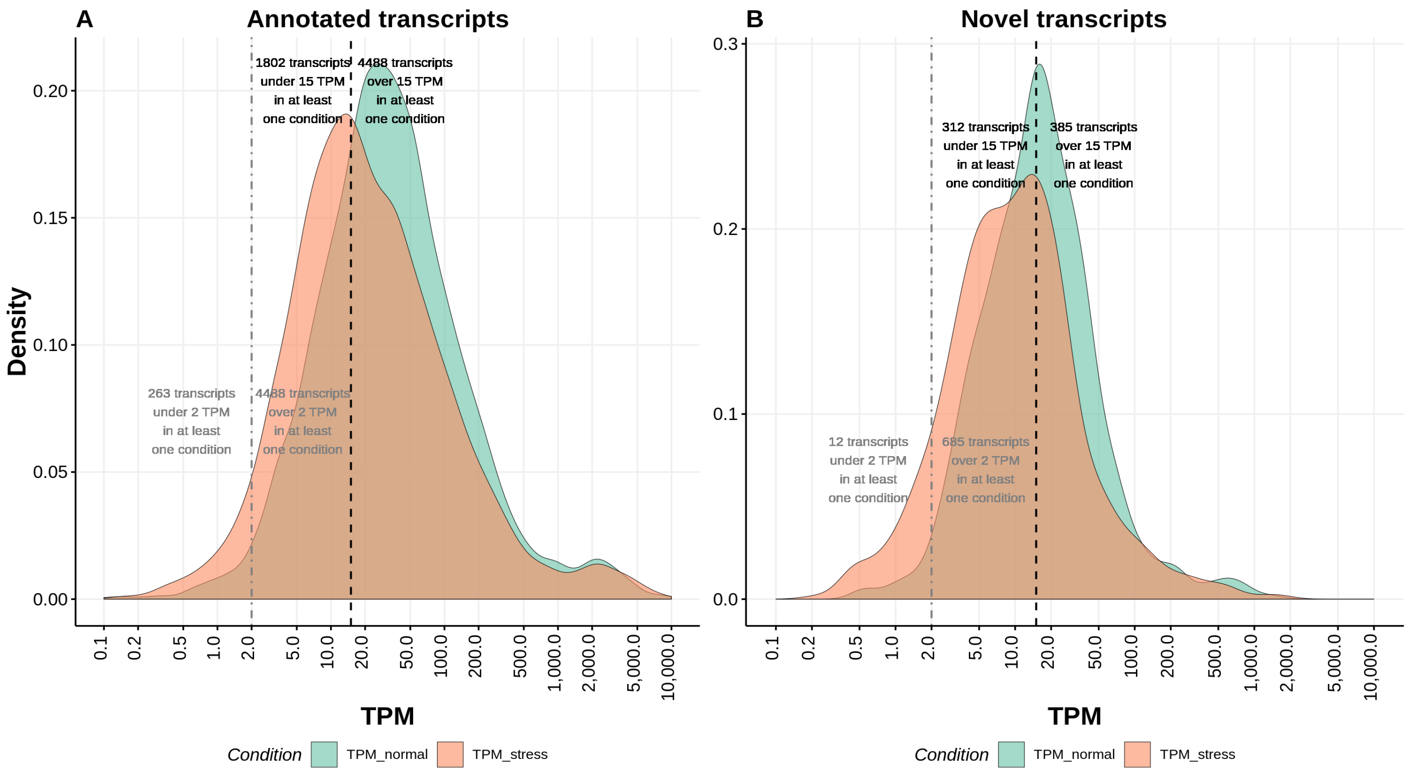


**Supplementary Figure 3 – Distribution of expression levels of annotated and novel transcripts. A.** Distribution of expression of annotated transcripts in rich media (green) and oxidative stress (orange) conditions. Grey line and text indicate a TPM cutoff of >2, black line and text indicate a TPM cutoff of >15. **B.** Distribution of expression of unannotated i.e. novel transcripts in rich media (green) and oxidative stress (orange) conditions. Grey line and text indicate a TPM cutoff of 2, black line and text indicate a TPM cutoff of 15.

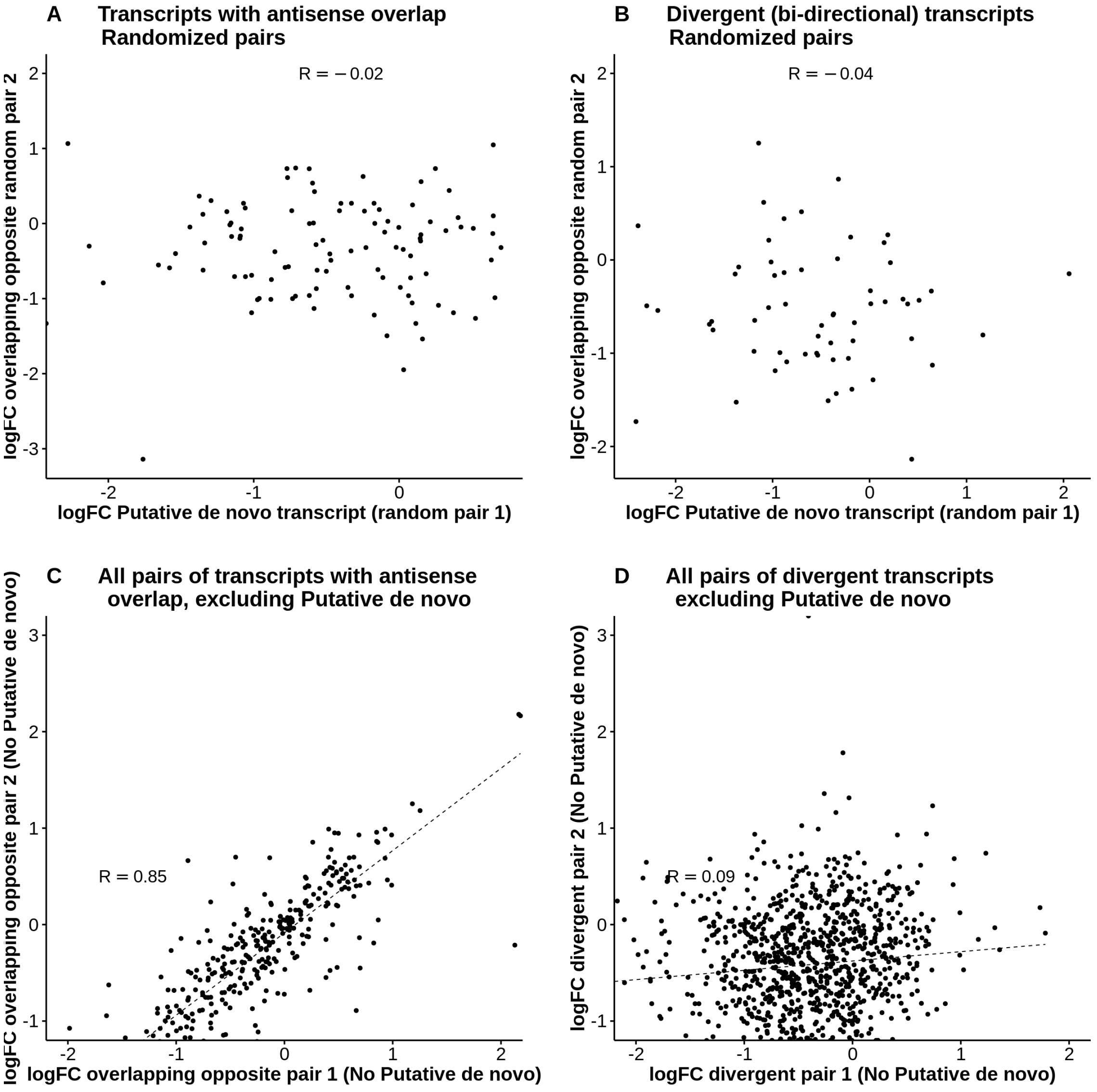


**Supplementary Figure 4. Randomized gene pairs do not show any significant correlation in expression values.** **A.** Log fold change (FC) of gene expression values in normal *versus* stress conditions for randomized pairs of overlapping antisense transcripts in which one of the pairs is a *de novo* transcript. Spearman's correlation -0.02 (p-value=0.8527). **B.** Log fold change (FC) of gene expression values in normal *versus* stress conditions for randomized pairs of divergent transcripts in which one of the pairs is a *de novo* transcript. The Spearman's correlation -0.04 (p-value=0.7392). **C.** Log fold change (FC) of gene expression values in normal *versus* stress conditions for pairs of overlapping antisense transcripts in which none of the pairs is a *de novo* transcript. Spearman's correlation 0.85 (p-value < $10^{-5}$). **D**. Log fold change (FC) of gene expression values in normal *versus* stress conditions for pairs of divergent transcripts in which none of the pairs is a *de novo* transcript. Spearman's correlation 0.09 (p-value < 0.0 0.006816).

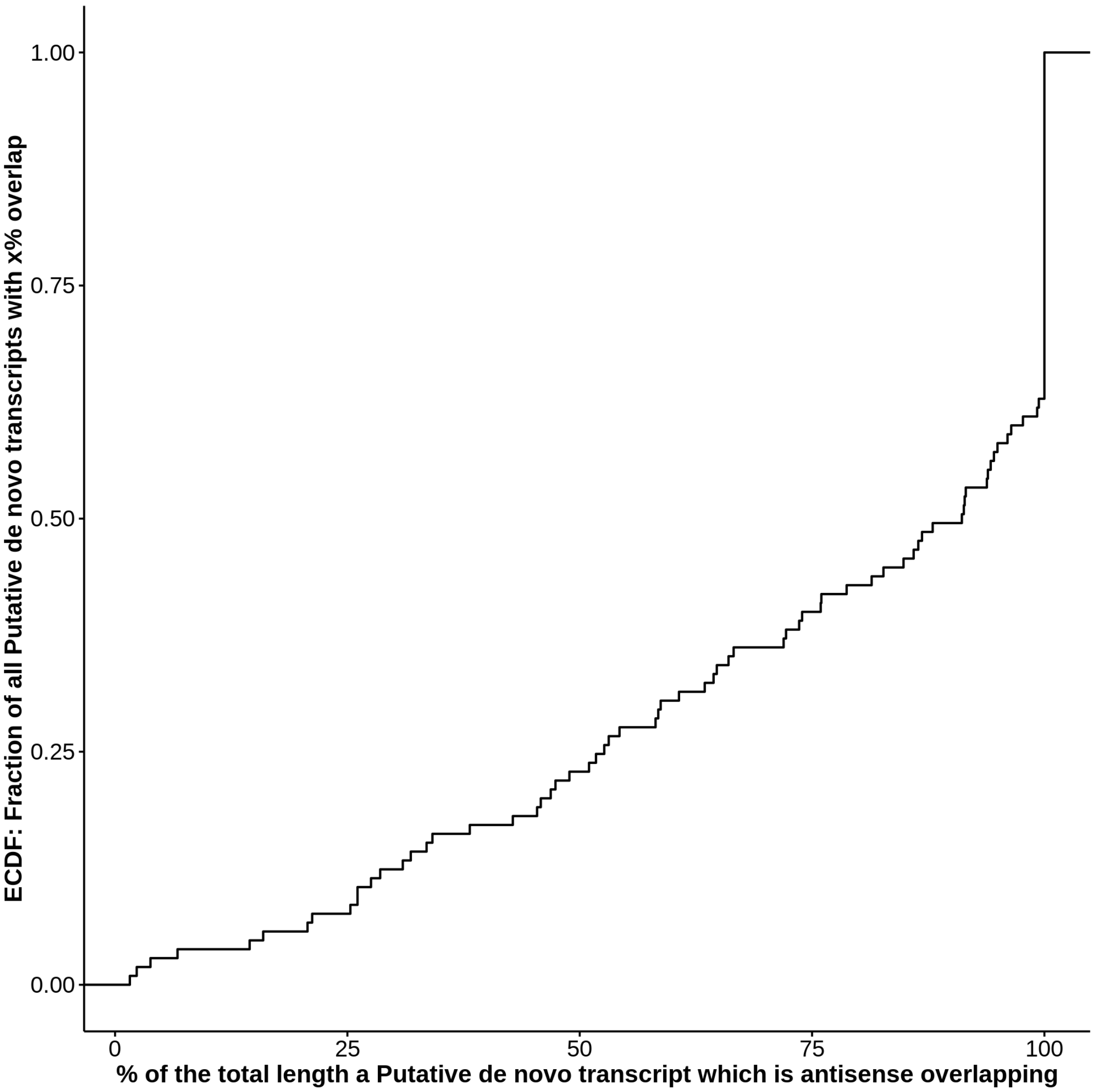


**Supplementary Figure 5. Putative *de novo* transcripts which overlap another transcript on the opposite strand tend have significant overlap.** This cumulative density function shows that the vast majority of putative *de novo* transcripts with any antisense overlap (minimum of 1 nt) have significant overlap with the other transcript. In half of these cases, the putative *de novo* transcript is overlapping for ~90% or more of their total length.

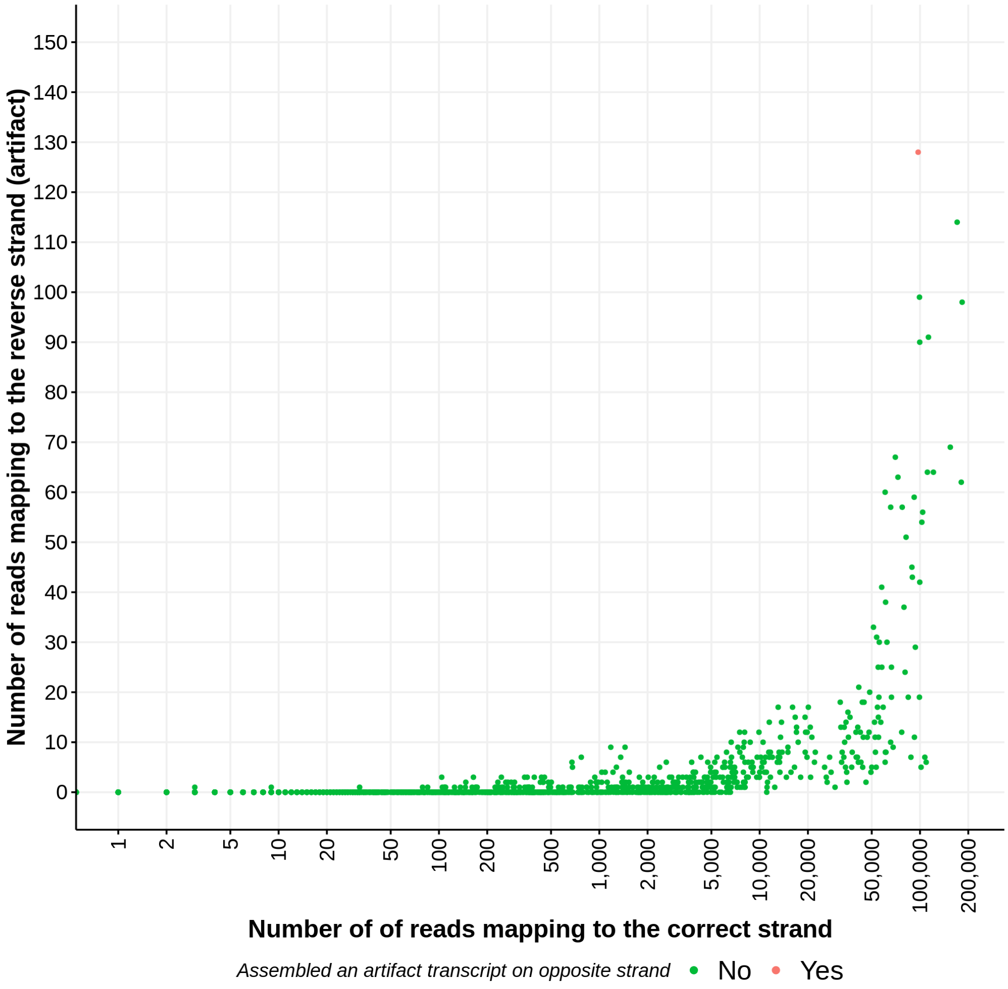


**Supplementary Figure 6. The fraction of reads that erroneously map to the antisense strand is negligible.** We included a spike-in (ERCC RNA spike-in mix 1) in each of our samples. The mix contains 92 synthetic transcripts at known concentrations. We recorded the number of reads that map to the sense and antisense strand of the ERCC RNAs. We observed that a tiny fraction of reads mapped in the opposite orientation; if there were enough of these artifacts, our pipeline could theoretically assemble a spurious novel transcript. We identified one case for which our pipeline assembled a spurious transcript; this occurred only for the most abundant spike-in transcript, ERCC-00130, which had a TPM over 66,000. However, this spurious transcript only appeared in one out of a possible 24 samples. In summary, we are confident that our novel transcripts which appear antisense overlapping to other transcripts are not spuriously assembled due to artifact reads which have mapped to the wrong strand.

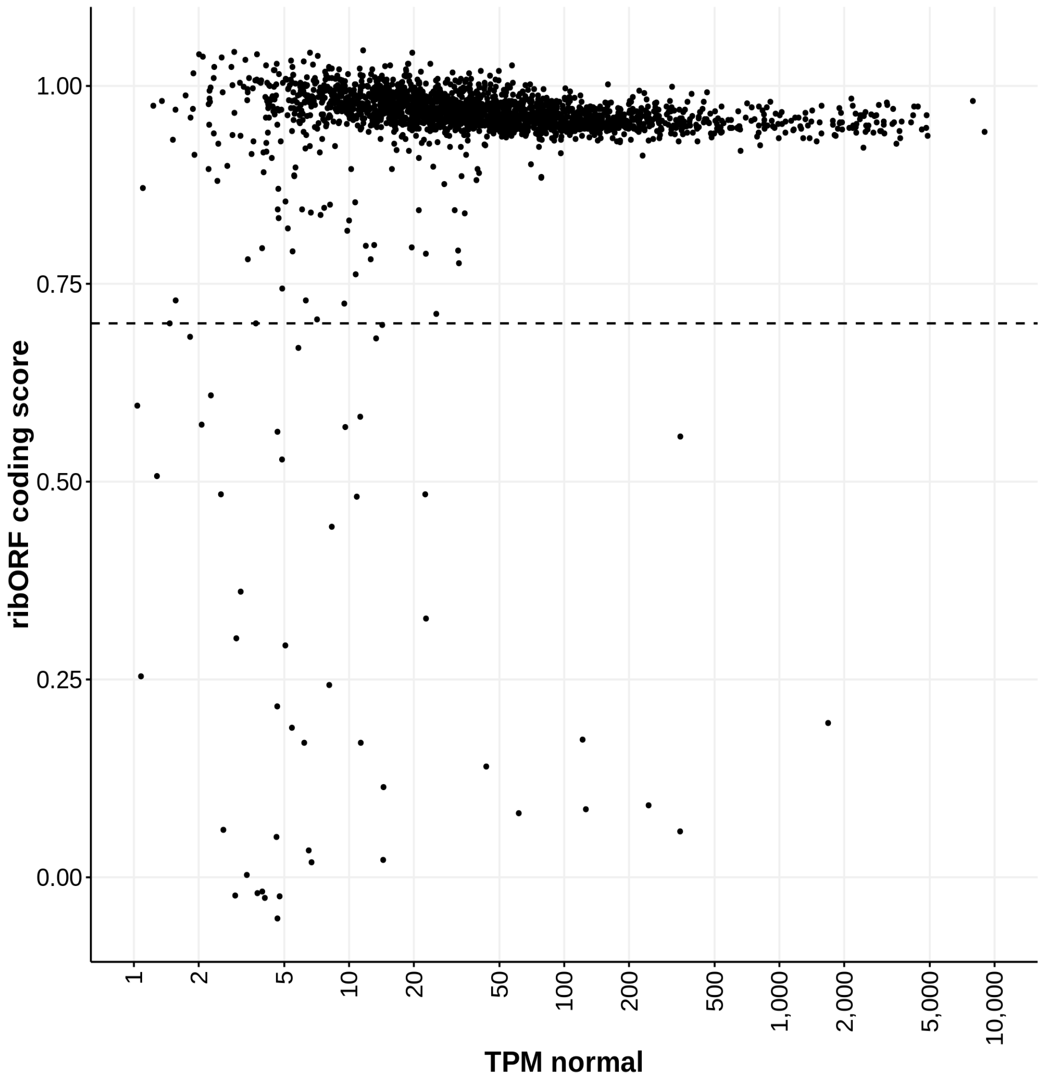


**Supplementary Figure 7. RibORF correctly classifies most verified annotated genes as coding when using our Ribo-Seq data.** We ran RibORF on a set of all verified ORFs from the *S. cerevisiae* reference annotations. 97.7% (2216/2270) of verified ORFs which had at least 10 mapped ribosome profiling reads were classified as coding by RibORF using a cutoff of 0.7, indicated by a dotted black line in the plot. In general, transcripts with higher expression were even more likely to be correctly classified as coding by RibORF.

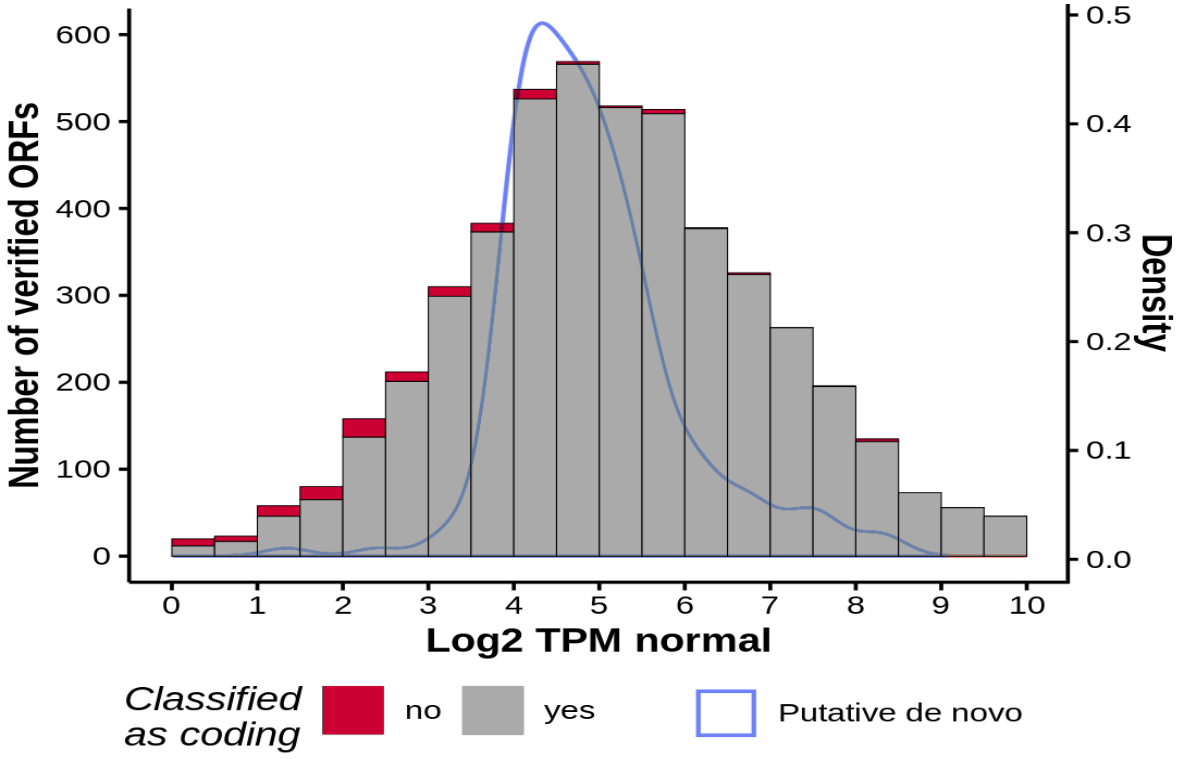


**Supplementary Figure 8. 2-5% of putative *de novo* genes may be incorrectly classified as non-coding by RibORF.** Based on the accuracy of RibORF's classification of verified annotated protein-coding (grey= correctly classified as translated, red= incorrectly classified as not translated) genes at different expression levels, we estimate that an additional 2-5% of putative *de novo* genes are incorrectly identified as non-coding by RibORF. This is due to the fact that RibORF performs best at higher expression levels (RNA-Seq) as there are typically more ribosome protected fragments to analyze; however, putative *de novo* transcripts tend to be expressed at lower levels than annotated coding genes (light blue line).

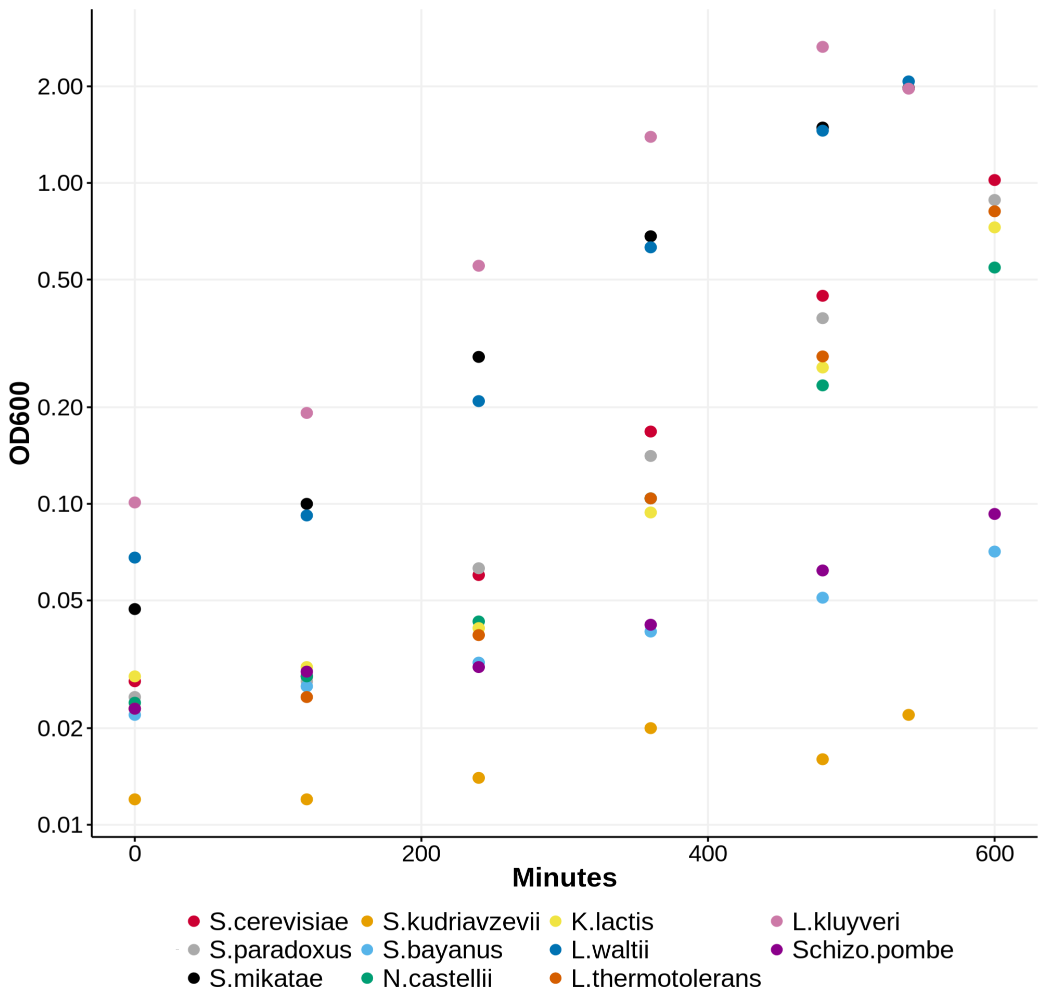


**Supplementary Figure 9. Many species of yeast grow well in the rich media developed by Tsankov et al. 2010.** Doubling times were calculated by recording a series of measurements of the $OD_{600}$ in rich media for all 11 species. Despite the differences in growth rate between species, we observed consistently shorter doubling times in the rich media than in YPD.

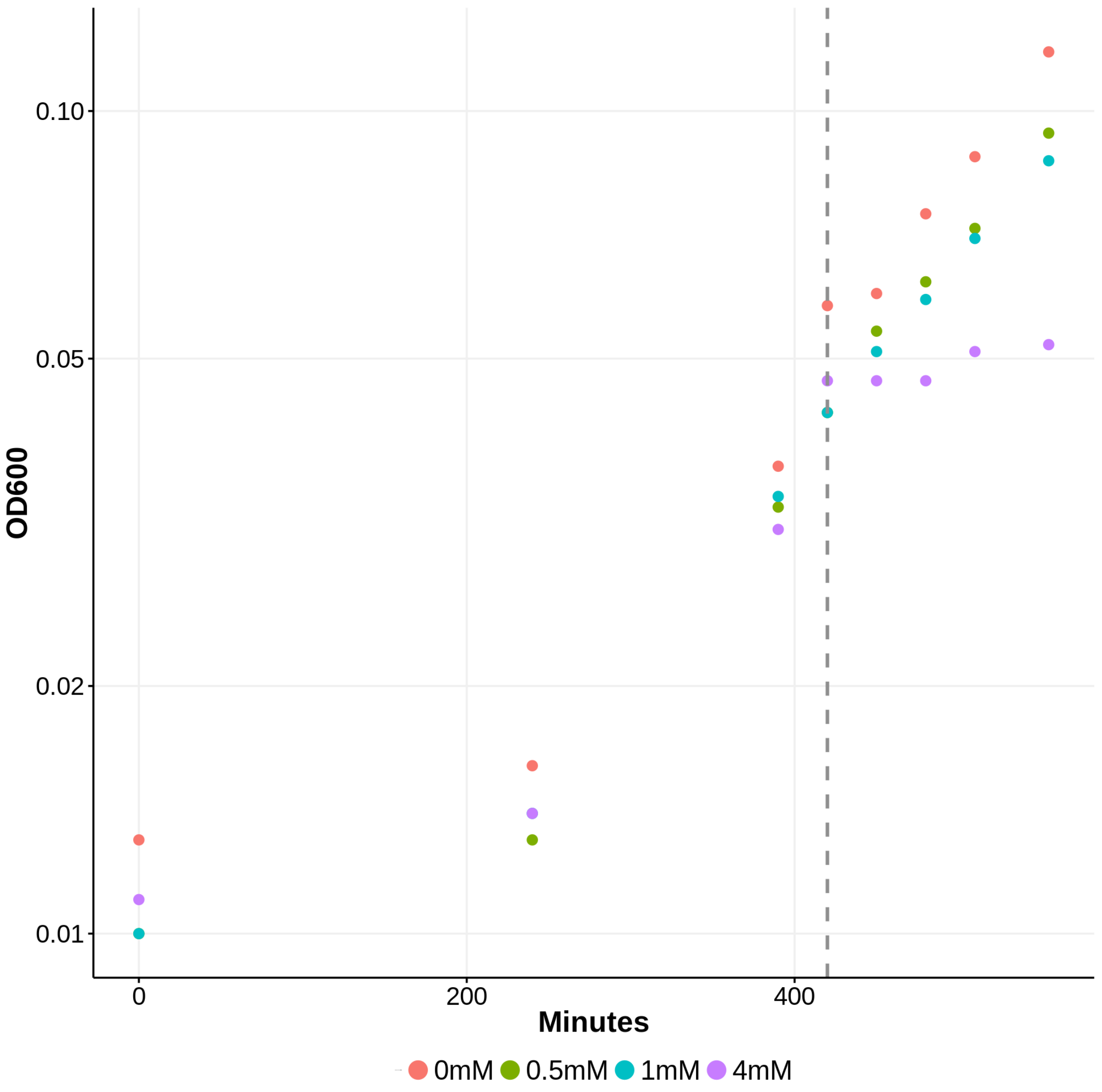


**Supplementary Figure 10. A concentration of 1.5mM $H_2O_2$ halves growth rate.** Growth rate curves were calculated for each species- in this case *N. castellii*- before and after the addition of $H_2O_2$. In this experiment the $H_2O_2$ was added to the rich media at minute 420 which is indicated on the plot by a dashed line. In the control condition 0mM $H_2O_2$ (in red) the yeast had a doubling time of 107 minutes. In the highest concentration $H_2O_2$ of 4mM, the yeast had a ~5x slower doubling time (543 minutes). The addition of a concentration of 1mM $H_2O_2$ resulted in a ~1.26x slower doubling time (135 minutes). Using 0.5 mM $H_2O_2$ increased the doubling time ~1.16x (124 minutes). We decided to use a final concentration of 1.5mM $H_2O_2$ for an approximate 2x slower doubling time i.e. halving the growth rate.
